# Impaired Experience-Dependent Theta Oscillation Synchronization and Inter-Areal Synaptic Connectivity in the Visual Cortex of Fmr1 KO Mice

**DOI:** 10.1101/2024.07.23.601989

**Authors:** Xi Cheng, Sanghamitra Nareddula, Hao-Cheng Gao, Yueyi Chen, Tiange Xiao, Yididiya Y. Nadew, Fan Xu, Paige Alyssa Edens, Christopher J. Quinn, Adam Kimbrough, Fang Huang, Alexander A. Chubykin

**Affiliations:** Department of Biological Sciences, Purdue Institute for Integrative Neuroscience, Purdue Autism Research Center, Purdue University, West Lafayette, IN, 47907, USA; Weldon School of Biomedical Engineering, Purdue University, West Lafayette, IN, USA; Department of Basic Medical Sciences, College of Veterinary Medicine, Purdue University, West Lafayette, IN 47906, USA; Department of Computer Sciences, Iowa State University, Ames, IA 50011, USA; School of Optics and Photonics, Beijing Institute of Technology, Beijing 100081, China; Advanced Research Institute of Multidisciplinary Science, Beijing Institute of Technology, Beijing 100081, China; Purdue Institute for Integrative Neuroscience, Institute for Cancer Research, Purdue University, West Lafayette, IN, USA; Purdue Institute of Inflammation, Immunology, and Infectious Disease, Purdue University, West Lafayette, IN, USA

## Abstract

Fragile X syndrome (FX) is the most prevalent inheritable form of autism spectrum disorder (ASD), characterized by hypersensitivity, difficulty in habituating to new sensory stimuli, and intellectual disability. Individuals with FX often experience visual perception and learning deficits. Visual experience leads to the emergence of the familiarity-evoked theta band oscillations in the primary visual cortex (V1) and the lateromedial area (LM) of mice. These theta oscillations in V1 and LM are synchronized with each other, providing a mechanism of sensory multi-areal binding. However, how this multi-areal binding and the corresponding theta oscillations are altered in FX is not known. Using iDISCO whole brain clearing with light-sheet microscopy, we quantified immediate early gene Fos expression in V1 and LM, identifying deficits in experience-dependent neural activity in FX mice. We performed simultaneous in vivo recordings with silicon probes in V1 and LM of awake mice and channelrhodopsin-2-assisted circuit mapping (CRACM) in acute brain slices to examine the neural activity and strength of long-range synaptic connections between V1 and LM in both wildtype (WT) and Fmr1 knockout (KO) mice, the model of FX, before and after visual experience. Our findings reveal synchronized familiarity-evoked theta oscillations in V1 and LM, the increased strength of V1→LM functional and synaptic connections, which correlated with the corresponding changes of presynaptic short-term plasticity in WT mice. The LM oscillations were attenuated in FX mice and correlated with impaired functional and synaptic connectivity and short-term plasticity in the feedforward (FF) V1→LM and feedback (FB) LM→V1 pathways. Finally, using 4Pi single-molecule localization microscopy (SMLM) in thick brain tissue, we identified experience-dependent changes in the density and shape of dendritic spines in layer 5 pyramidal cells of WT mice, which correlated with the functional synaptic measurements. Interestingly, there was an increased dendritic spine density and length in naïve FX mice that failed to respond to experience. Our study provides the first comprehensive characterization of the role of visual experience in triggering inter-areal neural synchrony and shaping synaptic connectivity in WT and FX mice.

## Introduction

Fragile X syndrome (FX) is the most common inherited form of autism spectrum disorders (ASDs) and intellectual disability^1^. Multiple studies have reported deficits in visual perception and learning in FX patients^2–5^. In a mouse model of FX, Fmr1 KO mice, sensory dysfunction, especially hypersensitivity, delayed learning on a visual discrimination task, and impaired perceptual learning have also been reported^6,7^. Previously, we found that visual familiarity is encoded by persistent theta (4-8 Hz) frequency oscillations in both single units and local field potentials (LFPs) in V1 of mice^8^. In Fmr1 KO mice, visual familiarity-evoked theta frequency oscillations were found to be attenuated, exhibiting lower power and shorter duration, suggesting deficits in encoding familiar stimuli in the primary visual cortex (V1)^9^. In addition to these impairments in visual perception and learning, multiple circuit and synaptic defects have been described in Fragile X patients and Fmr1 KO mice, including immature synaptic features^10^, increased density of unstable dendritic filopodia, and impaired experience-dependent and other types of synaptic plasticity, such as decreased long-term potentiation^11–13^. The abnormal short-term plasticity essential for information processing^14^, and working memory^15^, was also impaired in excitatory hippocampal synapses of Fmr1 KO mice^16^. However, the short-term potentiation (STP) alterations in the cortical regions of Fmr1 KO mice have not been characterized. An imbalance of neocortical excitation and inhibition was reported in Fmr1 KO mice, reflecting network hyperexcitability^17^, mediated by decreased activity of Parvalbumin positive (PV+) interneurons in the primary visual cortex (V1) of adult FX mice^6,18^. Consistent with these findings, we have observed impaired visual experience-dependent changes across inter-laminar synaptic connections in V1, particularly in projections from layer 5 (L5) pyramidal cells to layer 4 (L4) interneurons, mapped using Channelrhodopsin-2-assisted circuit mapping (CRACM) ^9^.

Recent decades of research have shown that visual processing in mice, like in primates, extends beyond V1 to higher-order visual areas (HVAs) that significantly influence V1 activity. The mouse visual cortex comprises multiple distinct areas, similar to primates, organized retinotopically and hierarchically, including anterolateral (AL), lateromedial (LM), and posteromedial (PM) areas^19–22^. V1 is interconnected with HVAs through feedforward (FF) and feedback (FB) pathways following the laminar projection patterns^23^ where FF connections transmit information from lower visual areas to HVAs, and FB operates in the opposite direction^24^. The LM in mice corresponds to the ventrotemporal areas in primates involved in object recognition, forming part of the ventral pathway^22,25,26^. In autism spectrum disorders (ASDs), an aberrant interplay between these bottom-up and top-down visual sensory processes may underlie the observed cognitive deficits^27^. Previous work in human patients reported reduced activity and connectivity in higher visual areas during tasks involving face processing, object recognition, and social perception. The fusiform face area is critical for face processing and demonstrates reduced activation and impaired functional connectivity with other brain areas involved in social cognition and emotional processing^28,29^.

To dissect the underlying mechanisms of the atypical visual perception and learning in FX, we performed simultaneous extracellular recordings of V1 and LM activity using silicon probes and to measure interareal connectivity by subcellular CRACM (sCRACM)^30^ between V1 and LM in both naïve and visually experienced WT and Fmr1 KO mice. Here, we report that the 4-8 Hz visual familiarity elicited oscillations of both LFP and single-unit population activity were lower in power and shorter in duration, and unit population firing rates were lower in V1 and LM of Fmr1 KO mice compared to WT mice. These alterations in neural activity were also verified by c-Fos quantification, an immediate early gene product used as a cell activation marker, by individual neurons in V1 and LM^31^. We measured the balance of excitation and inhibition, as well as STP, in FF V1→LM and FB LM→V1 pathways utilizing CRACM and optogenetic stimulated paired-pulse, respectively. Visual experience-dependent changes in synaptic strength were impaired in Fmr1 KO mice, particularly at V1 to LM L2/3 and L5 PCs connection in FF V1→LM pathway and LM to V1 L5 connection in FB LM→V1 pathway. The deficits in dendritic spine plasticity, characterized by thinner spine necks and unchanged spine head diameters after visual experience, were detected in Fmr1 KO mice by 4Pi single-molecule localization microscopy ^32^. These morphological alterations could underlie the observed synaptic strength deficits in Fmr1 KO mice. Furthermore, the balance between excitation and inhibition, along with STP, was impaired in FF V1→LM and FB LM→V1 pathways in Fmr1 KO mice. Altogether, our findings highlight deficits in communication and impaired synaptic plasticity between LM and V1 in Fmr1 KO mice.

## Results

### Whole brain clearing revealed impaired c-fos activity following visual experience in V1 and LM of FX mice

Our previous work has demonstrated the emergence and synchronization of oscillations in higher visual areas with those in V1 ^33^. The immediate early gene (IEG) c-Fos is a well-established marker of neuronal activity and plasticity, making it an ideal target for studying experience-dependent changes in the brain ^34^. We sought to elucidate how visual stimuli influence neuronal activity and plasticity, particularly in conditions associated with neural hyperactivity, such as Fragile X syndrome (FX) by analyzing Fos expression in mouse visual areas before and after visual experience. We subjected mice to a passive visual perception paradigm to investigate the difference in the neural activity evoked by visual perceptual experience in V1 and LM of FX mice. To establish visual response before visual experience, mice were habituated to the experimental setup for 2-3 days and then trained to a filtered pink noise stimulus (0.12 cycles/degree of visual angle (cpd), 0.75 Hz) corresponding to the spatial frequency (SF) and time-frequency (TF) shown to elicit the strongest response in LM^35^. The training was continued for 4-6 days, wherein the filtered pink noise movie was presented for 200 ms, 200 times a day, with a grey screen between trials (6 s ITI, Figure 1A). We decided to investigate how perceptual experience alters Fos expression in the visual cortical areas, specifically in V1 and LM, using a combination of the iDISCO brain clearing technique and light sheet microscopy for fast volume imaging of whole-brain samples (Figure 1B, C).

**Figure 1.**
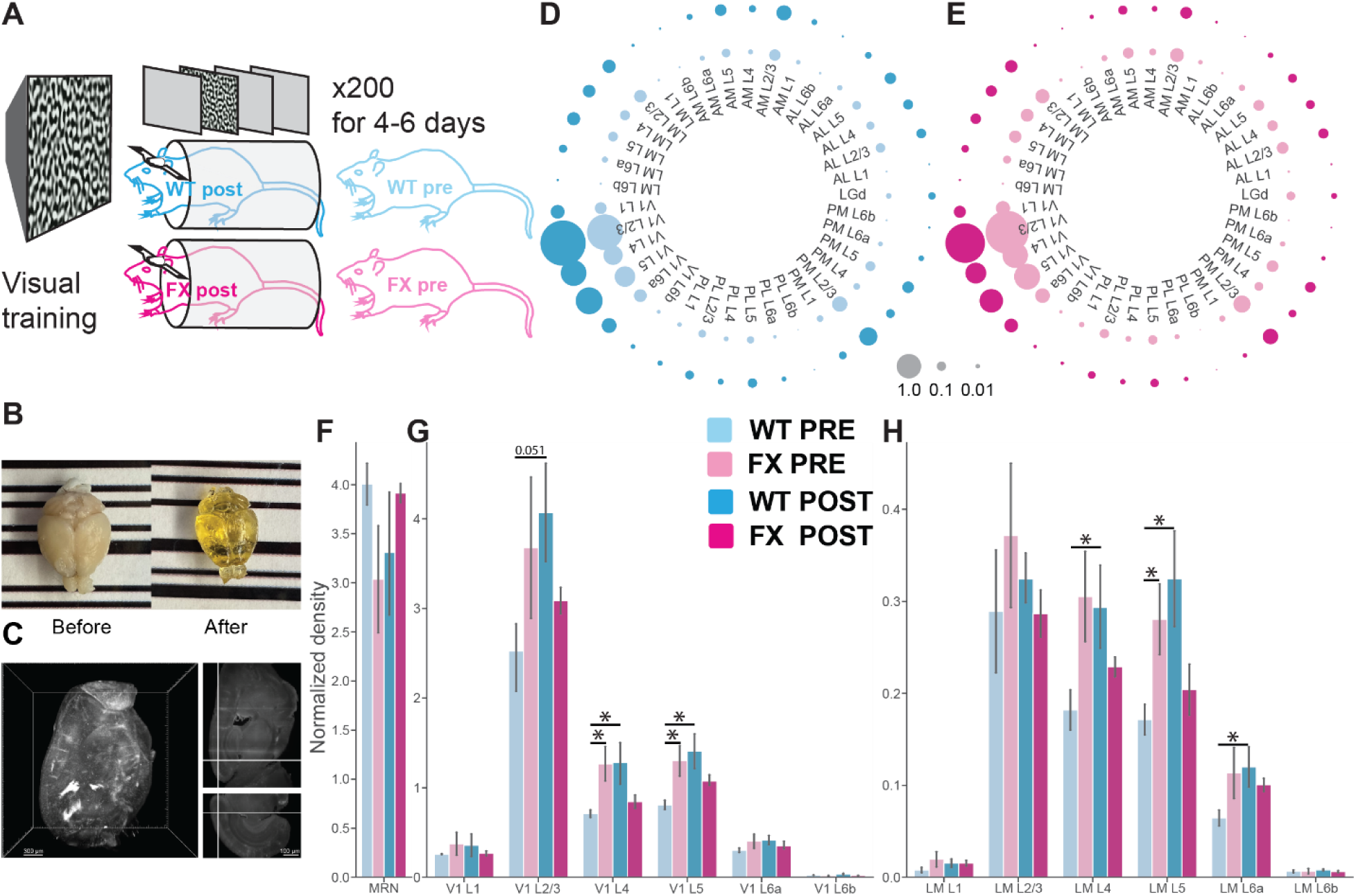
Normalized C-fos+ expression in V1 and LM. **(A)** Schematic representation of visual familiarity paradigm for WT and FX animals. **(B)** The whole mouse brain before and after iDISCO clearing. **(C)** Projection of a whole-mount c-Fos immunolabeling of a mouse brain in the sagittal direction, shown V1 and HVA (Left). The example of c-Fos immunolabeling in a sagittal (top right) and coronal cross-section (bottom right). **(D)** C-fos level (normalized density) in different visual areas represented by the circle size in WT mice. **(E)** Same as D) but in FX mice. **(F)** Bar graphs of average c-fos+ cell number in MRN normalized to overall c-fos number (averaged by all available brain regions) in a mouse (WT pre: N=4, WT post: N=4, FX pre: N=4, FX post: N=4 animals). The significance was reported by the T-test (p values are listed in Table X). **(G)** Bar graphs of average c-fos+ cell number in V1 normalized to it in MRN. The significance was reported by the T-test. **(H)** Bar graphs of average c-fos+ cell number in LM normalized to it in MRN. The significance was reported by the T-test. Data were presented as mean ± SEM. T-test: *p<0.05.

We measured Fos expression in four groups of animals: WT naive, FX naive, WT post-experience, and FX post-experience. To normalize the quantification of Fos expression across different mice, we used the median raphe nucleus (MRN), a midbrain region known to regulate many essential functions in mice, as a control. As expected, we found no difference in Fos expression from neurons in MRN among the four groups of mice (Figure 1F). Fos expression was generally much higher in V1 than in the other visual areas, including LM (Figure 1D, E, Figure S1). Visual perceptual experience increased in Fos expression in multiple layers of V1, including L2/3, L4, L5, and in L4, L5, and L6a of LM in WT, but FX mice (Figure 1G, H). Interestingly, the number of Fos-positive neurons was higher in FX mice before visual experience in multiple layers of V1: L4, L5, LM L5 (Figure 1G, H), LGN, AM: L4, L5, L6b, PL L6a, and PM: L4, L5 (Figure S1). These results are consistent with the previous reports of cortical hyperactivity in visual areas of FX mice. Overall, visual experience induced a prominent upregulation in Fos-positive neuron density in multiple visual areas of WT mice, suggesting induction of experience-dependent neuronal and/or synaptic plasticity. The changes in Fos density between pre- and post-visual experience in FX mice did not reach significance across layers in both V1 and LM, suggesting impaired visual experience-dependent plasticity.

### The attenuated visual experience evoked oscillations and synchrony in LFPs of FX mice in V1 and LM

To determine how visual experience influences dynamic interaction and sensory binding between visual areas, we decided to perform simultaneous extracellular recordings from V1 and LM before and after visual experience. We applied the same visual experience paradigm as in the Fos experiment, activating both V1 and LM to WT and FX mice. 64-channel silicon probes were inserted normally into V1 and LM to record simultaneously from both regions after the habituation of the mice to the experimental setup and after the visual experience (Figure 2A, B). LFPs were evaluated by selecting the channel with the strongest activity post visual stimulus presentation (0.5-1.0 s) averaged across 20 trials, normalized to baseline activity (0-0.5 s). In both V1 and LM visual experience induced strong stimulus-locked theta oscillatory activity (4-8 Hz), persisting past the visual stimulus offset (Figure 2C). We evaluated these oscillations by quantifying the amplitude and power of the LFP activity. Post visual experience VEP amplitudes were significantly larger in magnitude for the second peak after stimulus onset for both V1 and LM in WT mice. However, FX mice showed no significant increase in VEP amplitudes post visual experience compared to naïve mice (Figure 2E, F). This attenuation of oscillations in FX mice is consistent with our previous reports in V1 ^9,36^. To characterize the magnitude of oscillatory activity in LM of WT and FX mice, performed time-frequency analysis of VEPs across several ranges of frequencies (4-8 Hz, 8-12 Hz, 12-30 Hz, 30-40 Hz, 50-70 Hz, and 30-70 Hz) (Figure 2I-L). Power spectrum analysis of the integral averaged power values within each frequency band showed a significant loss in power within the 4-8 Hz range of FX mice compared to WT mice in both V1 and LM post-visual experience (Figure 2K, L). These observations could indicate impaired inter-areal communication between V1 and LM in encoding visual familiarity in FX mice. To quantify the synchrony between V1 and LM, we computed the phase difference between 4-8 Hz oscillations in V1 and LM and discovered a decrease in phase difference between V1 and LM for 4-8 Hz oscillations post-visual experience in WT mice, indicating a strengthening of synchronization between the two regions (Figure 2G, H). However, this phase synchronization was not seen in FX mice, further indicating a decrease in synchrony and dynamic binding between V1 and LM in FX mice (Figure 2G, H).

**Figure 2.**
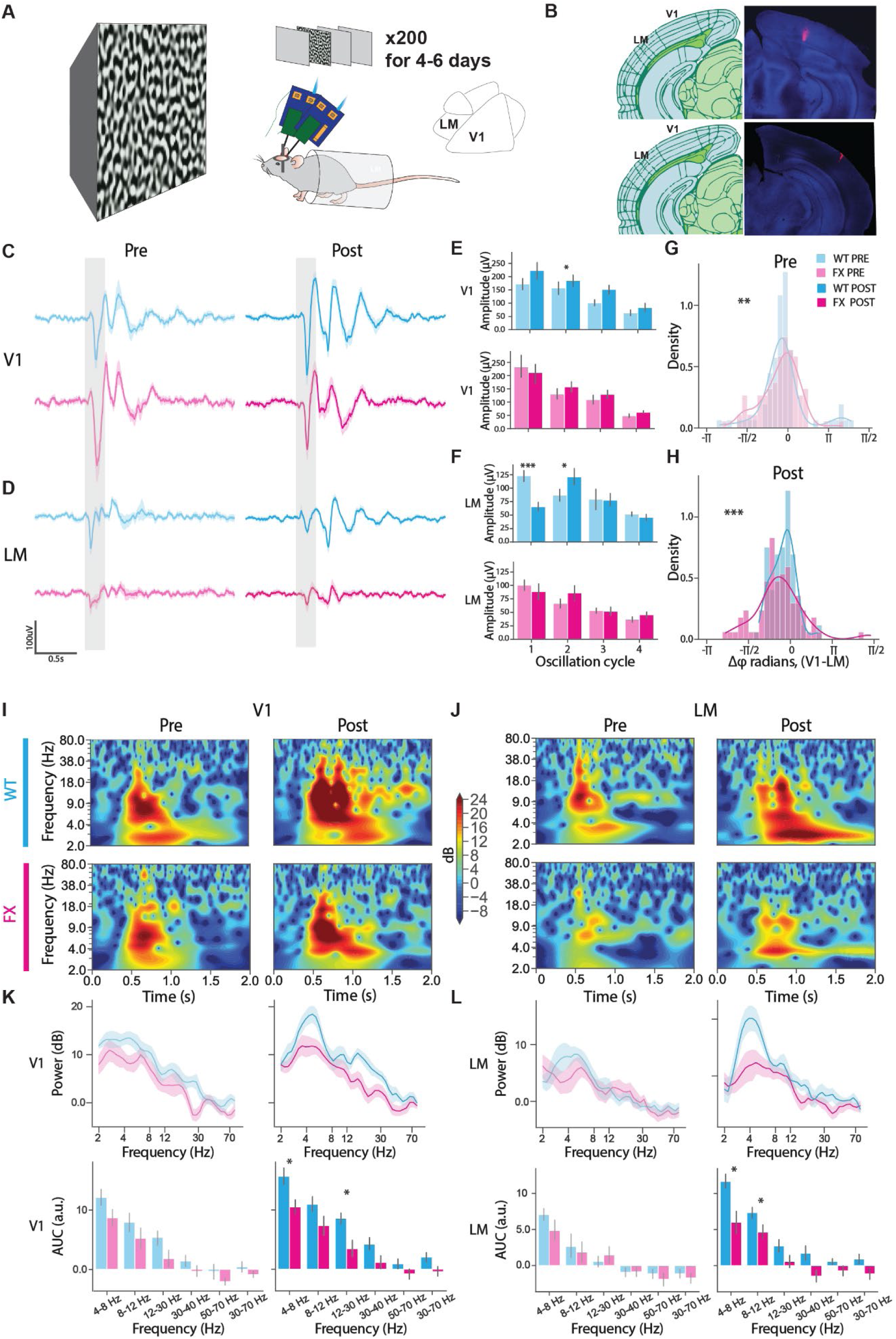
LFPs from simultaneous recordings in V1 and LM show attenuation in visual experience-evoked oscillatory activity in FX mice compared to WT mice. **(A)** Schematic representation of simultaneous LM-V1 recordings in awake head-fixed mice. **(B)** Representative histology showing probe traces of recording sites for V1 and LM. Probes were dipped in DiD Vybrant cell labeling dye and inserted at the recording locations. **(C)** Averaged visually evoked potentials (VEPs) in V1 of WT (cyan) and FX (magenta) mice. Trial averaged VEPs in layer 4 of V1, pre (left; WT: n=10 mice, FX: n=8 mice) and post (right; WT: n=11 mice, FX: n=9 mice) visual experience. **(D)** Averaged visually evoked potentials (VEPs) in LM of WT (cyan) and FX (magenta) mice. Trial averaged VEPs in LM, pre (left; WT: n=10 mice, FX: n=8 mice) and post (right; WT: n=11 mice, FX: n=9 mice) visual experience. **(E)** V1 VEP amplitudes at 4 oscillation cycles averaged across mice, Top: WT mice (cyan) (Pre: n=10 mice, Post: n=11 mice); Bottom: FX mice (magenta) (Pre: n=8 mice, Post: n=9 mice). Two-way ANOVA followed by Tukey HSD; V1 WT; Oscillation cycle 2: p=0.04. **(F)** LM VEP amplitudes at 4 oscillation cycles averaged across mice, Top: WT mice (cyan) (Pre: n=10 mice, Post: n=11 mice); Bottom: FX mice (magenta) (Pre: n=8 mice, Post: n=9 mice). Two-way ANOVA followed by Tukey HSD; LM WT; Oscillation cycle 1: p=0.0001, Oscillation cycle 2: p=0.042. **(G)** Density plots showing phase difference between V1 and LM 4-8 Hz oscillations after stimulus onset (0.5-2s), pre training. Two-sample Kuiper test; Pre training; D=0.315, p=3.65 x 10^-3^. **(H)** Density plots showing phase difference between V1 and LM 4-8 Hz oscillations after stimulus onset (0.5-2s), post training. Two-sample Kuiper test; Post training; D=0.341, p=8.33 x 10^-4^. **(I)** Heat maps showing time-frequency spectrograms of V1 trial averaged VEPs in WT (cyan) and FX (magenta) mice. Top: Time-frequency spectrograms of layer 4 V1 trial averaged VEPs in WT mice pre and post training (Left; Pre: n=10 mice, Right; Post: n=11 mice). Bottom: Time-frequency spectrograms of layer 4 V1 trial averaged VEPs in FX mice pre and post training, pre (Left; Pre: n=8 mice, Right; Post: n=9 mice). **(J)** Heat maps showing time-frequency spectrograms of LM trial averaged VEPs in WT (cyan) and FX (magenta) mice. Top: Time-frequency spectrograms of LM trial averaged VEPs in WT mice pre and post training (Left; Pre: n=10 mice, Right; Post: n=11 mice). Bottom: Time-frequency spectrograms of LM trial averaged VEPs in FX mice pre and post training (Left; Pre: n=8 mice, Post: n=9 mice). **(K)** Quantified power across various frequency bands averaged within a time window of 0.7-1.25s of stimulus onset for trial averaged VEPs in WT (cyan) and FX (magenta) mice in V1. Top: Power spectrum across various frequency bands for trial averaged layer 4 V1 VEPs, pre (Left; WT: n=9 mice, FX: n=8 mice) and post (Right; WT: n=8 mice, FX: n=11 mice) visual experience. Area under the curve within various frequency bands for trial averaged layer 4 V1 VEPs, pre (Left; WT: n=9 mice, FX: n=8 mice) and post (Right; WT: n=8 mice, FX: n=11 mice) visual experience. Mann-Whitney U test; Post training; 4-8Hz: p=0.041, 12-30Hz: p=0.041. **(L)** Quantified power across various frequency bands averaged within a time window of 0.7-1.25s of stimulus onset for trial averaged VEPs in WT (cyan) and FX (magenta) mice in LM. Top: Power spectrum across various frequency bands for trial averaged LM VEPs, pre (Left; WT: n=9 mice, FX: n=8 mice) and post (Right; WT: n=8 mice, FX: n=11 mice) visual experience. Bottom: Area under the curve within various frequency bands for trial averaged LM VEPs, pre (Left; WT: n=9 mice, FX: n=8 mice) and post (Right; WT: n=8 mice, FX: n=11 mice) visual experience. Mann-Whitney U test; Post training; 4-8Hz: p=0.041, 8-12Hz: p=0.033.

### Unit population firing rates showed attenuation of visual experience evoked oscillations in LM and V1 of FX mice

To investigate the difference in unit population response between FX and WT mice, we analyzed baseline (0-0.5 s) normalized Z-score firing rates of units from V1 and LM. Unit population firing rates show persistent oscillatory activity in the 4-8 Hz theta range post-visual experience as seen previously ^9,36^ (Figure 3A, C, D). Prior to visual experience, unit population activity showed a single peak temporally locked to visual stimulus presentation in both V1 and LM. The visual experience evoked multiple oscillation peaks persisting long after stimulus presentation in V1 and LM. On average, WT mice typically showed 4 peaks or oscillation cycles, while FX mice showed fewer oscillation cycles (Figure 3D, E). Duration analysis of the detected peaks showed that this persistent oscillatory unit activity was shorter in V1 and LM of FX mice (Figure 3E). Mean persistent activity durations of WT mice post-visual experience were significantly longer than FX mice for both V1 and LM (Figure 3E). The maximum firing rate Z-scores were calculated for each unit within each oscillation cycle window. Average firing rates across the unit population were significantly higher for WT mice compared to FX mice for oscillation cycles following the initial peak in V1 (cycle 2, cycle 3, and cycle 4) and LM (cycle 2 and cycle 4), indicating an overall decrease in synchronous population firing of units (Figure 3F). The attenuated power of the theta frequency unit activity in FX mice is consistent with the reduced Fos activity in V1 and LM areas following visual experience.

**Figure 3.**
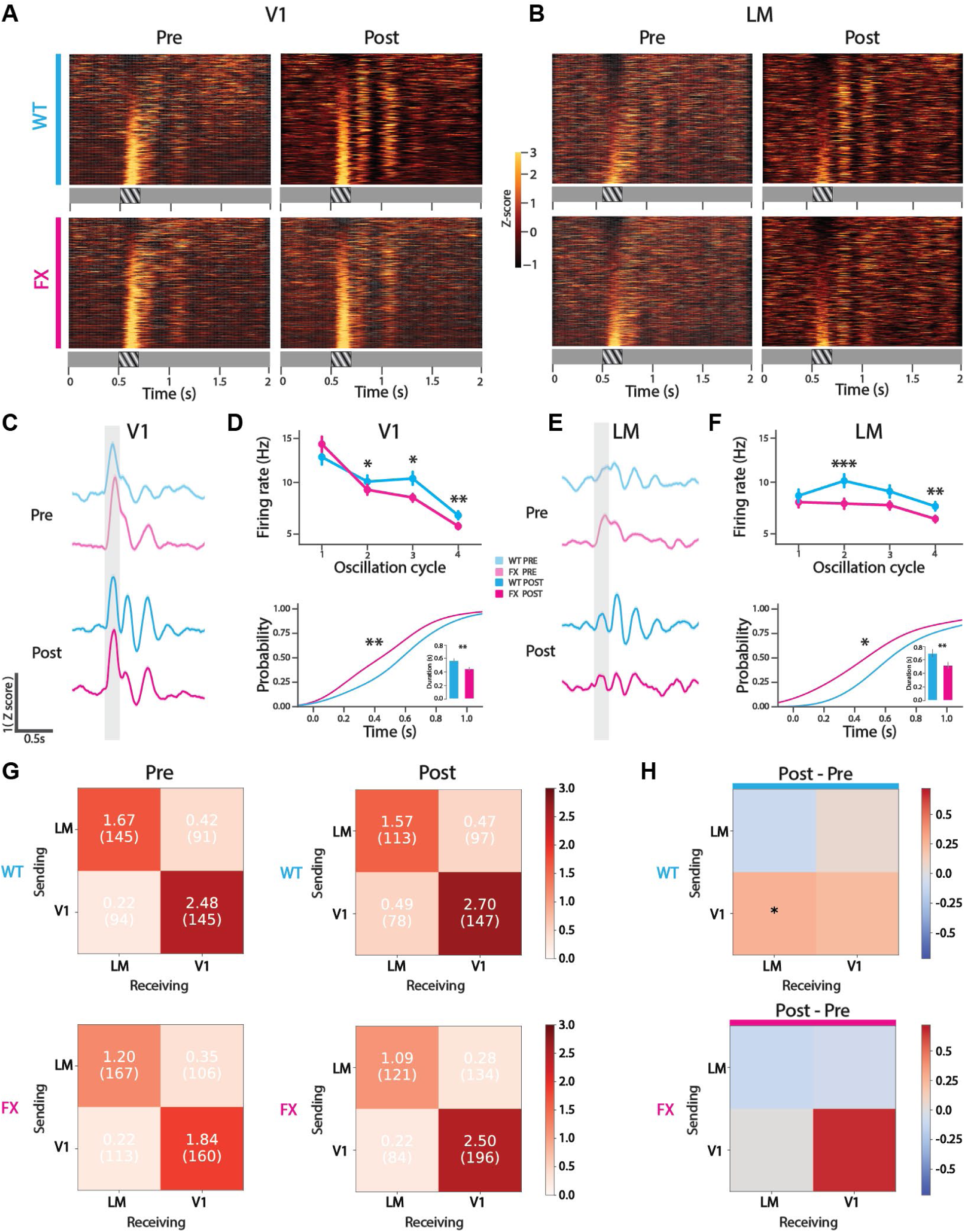
Unit population activity shows attenuation in visual experience evoked oscillations in V1 and LM of FX mice compared to WT mice. **(A)** Heat maps showing z-score firing rates of single units in V1, pre (left; WT: N=267 units, n=10 mice, FX: N=270 units, n=8 mice) and post (right; WT: N=224 units, n=11 mice, FX: N=327 units, n=9 mice) visual experience in WT (top, cyan) and FX (bottom, magenta) mice. **(B)** Heat maps showing z-score firing rates of single units in LM, pre (left; WT: N=322 units, n=10 mice, FX: N=410 units, n=8 mice) and post (right; WT: N=259 units, n=11 mice, FX: N=312 units, n=9 mice) visual experience in WT (top, cyan) and FX (bottom, magenta) mice. **(C)** Baseline normalized population averaged z-score firing rates of V1 units shown in **(A)**. **(D)** Top: Maximum firing rates at 4 oscillation cycle time windows averaged across V1 units shown in **(A)** post-training in WT (cyan) and FX (magenta) mice. Mann-Whitney U test; Cycle2: p=0.04, Cycle3: p=0.039, Cycle4: p=0.004 (WT: N= 224 units across 11 mice, FX: N= 327 units across 9 mice). Bottom: Cumulative distribution function (CDF) of the duration of identified local maxima of averaged z-score firing rates of V1 units shown in **(A)** post-visual experience (two-sample Kolmogorov-Smirnov test WT vs. FX, D=0.23, p=0.002). Inset: Mean peak duration post visual experience WT vs FX (Tukey HSD multiple comparison of means; p=0.0008) (WT: N= 129 units post peak detection across 11 mice, FX: N= 145 units post peak detection across nine mice). **(E)** Baseline normalized population averaged z-score firing rates of LM units, as shown in **(B)**. **(F)** Top: : Maximum firing rates at four oscillation cycle time windows averaged across LM units shown in **(B)** post-training in WT (cyan) and FX (magenta) mice. Mann-Whitney U test; Cycle2: p=2.24E-05, Cycle4: p=0.004 (WT: N= 259 units across 11 mice, FX: N= 312 units across 9 mice). Bottom: Cumulative distribution function (CDF) of the duration of identified local maxima of averaged z-score firing rates of LM units shown in **(B)** post-visual experience (two-sample Kolmogorov-Smirnov test WT vs FX, D=0.31, p=0.033). Inset: Mean peak duration post visual experience WT vs FX (Tukey HSD multiple comparison of means; p=0.019) (WT: N= 61 units post peak detection across 11 mice, FX: N= 101 units post peak detection across nine mice). **(G)** Directed information analysis of units across all layers of V1 and LM (Markov order of 10ms). Heat maps indicate the strength of connectivity, with darker colors representing stronger predictive connections. Top: Pre-training (left) and post-training (right) heat maps for WT mice. Bottom: Pre-training (left) and post-training (right) heat maps for FX mice. **(H)** Heat maps showing the difference between pre- and post-training connectivity changes using directed information analysis shown in **(G)** for WT (top) and FX (bottom) mice. *p<0.05.

### Directed information analysis revealed increased functional feedforward connectivity from V1 to LM in WT, but not FX mice following visual experience

We conducted a directed information analysis ^9,37,38^ to assess changes in putative functional connectivity between units in V1 and LM pre and post-training and to assess how those changes differed between WT and FX mice. The analysis entailed fitting parametric models to each unit’s spike train data conditioned on spike train data of other units, evaluating directed information on those unit-level models, and then aggregating unit-level values to estimate area-level connection strengths. This was repeated for each experimental setting (WT/FX, pre/post training). Differences in area-level connection strengths between different experimental settings were then calculated. We discovered a significant increase in the FF connectivity from V1 to LM post-visual experience in WT mice. However, in FX mice, no significant change was detected in this FF connectivity post-visual experience (Figure 3H). Additionally, the FB input from LM to V1 demonstrated a trend toward an increase in the strength of connectivity post-visual experience in WT mice, though it did not reach significance, whereas FX mice did not show this trend (Figure 3H). These changes in the functional connectivity following visual experience suggest that there may be changes in the strength of synaptic connections between V1 and LM after visual experience, which may also be altered in FX mice.

### The visual experience led to a strengthening of synaptic connectivity of FF V1→LM pathway from V1 to LM in WT mice, which was attenuated in FX Mice

To validate our in vivo directed information results, we decided to directly measure the visual experience-dependent synaptic plasticity of the FF V1→LM or FB LM→V1 projections in FX mice and WT mice using channelrhodopsin-assisted circuit mapping (CRACM) in acute brain slices ^9,39^. We first injected AAV1-ChR2-Venus into V1. Three weeks later, we subjected the mice to a passive visual perceptual experience for 4-6 days using the same paradigm we employed for in vivo experiments and c-fos labeling (Figure 4A). CRACM recordings from experienced mice were conducted the day after the end of the last round of perceptual training. WT and FX mice were pseudo-randomly assigned to either pre-training or experienced groups to avoid observation bias. The genotypes of mice were blinded to operators during the experiment and statistical analysis. To ensure that the expression levels of ChR2 did not affect synaptic strength measurements, the fluorescent intensity of the recording regions from each mouse of either V1 or LM injections were quantified and confirmed to be no different among groups (Figure S3). Fluorescent dye was added to the tip of the recording electrode with the internal solution for tracing the cell morphology in layers L2/3, L4, and L5 of both regions, and the sCRACM map was overlaid with the confocal imaging of the corresponding cells (Figure S4). The neurons patched in V1 were mostly principal cells (PCs) according to their morphological characteristics (Figure S4). The whole-cell patch clamp recordings of LM L2/3, L4, and L5 neurons were conducted in acute brain slices from four groups of animals: WT naive, FX naive, WT post-experience, and FX post-experience (Figure 4B, G, L).

**Figure 4.**
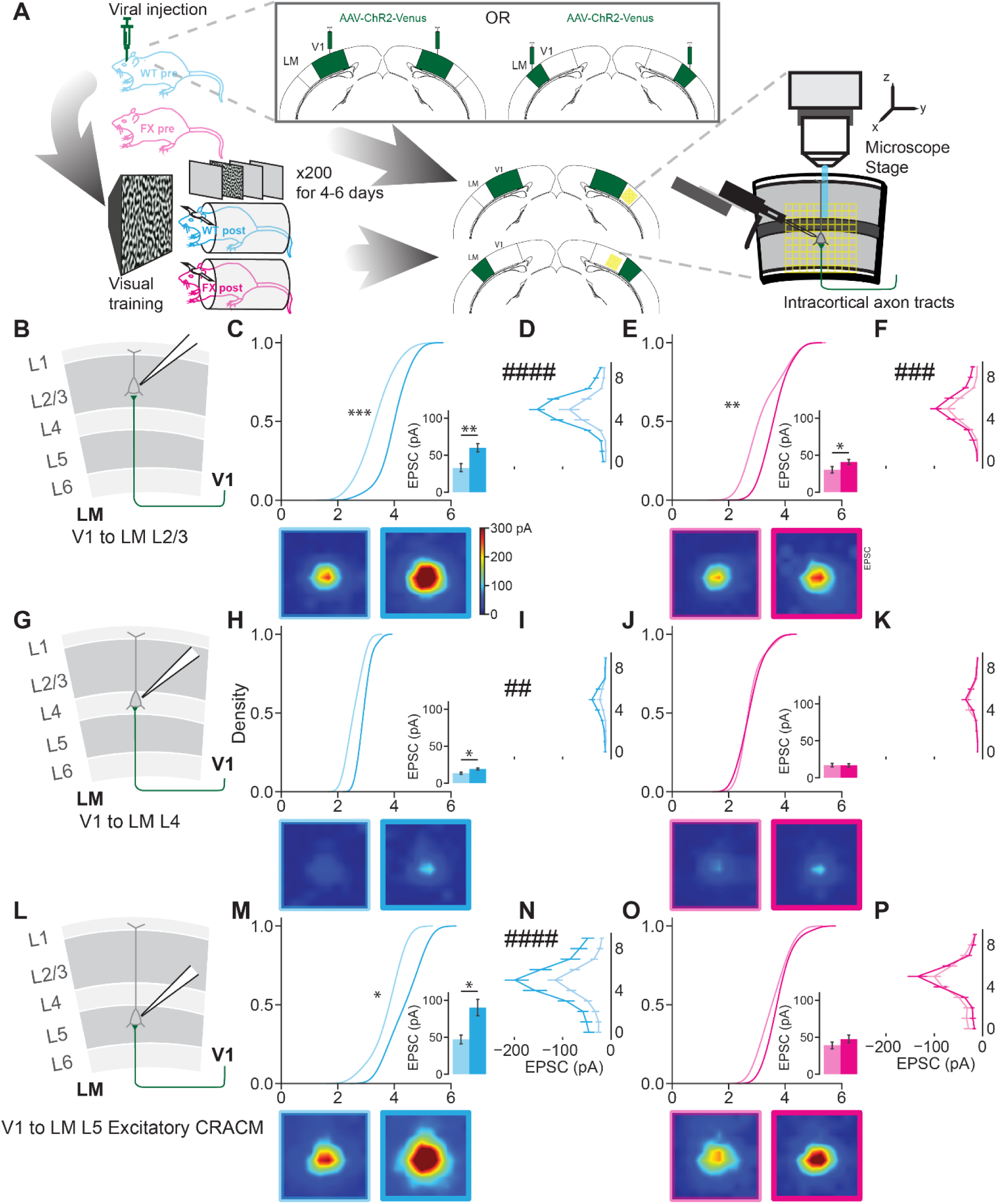
FF sCRACM maps of input from V1 ChR2 positive neurons to PCs in LM comparing between naïve and post perceptual experience groups. **(A)** Experimental workflow of CRACM experiment on acute brain slices from pre- and post- training mice. **(B)** Illustration of V1 to LM L23 PC projections. **(C)** The cumulative distributions (top) of the averaged EPSCCRACM amplitudes plotted in ln scale in sCRACM map recorded from LM L23 PCs. Significance was reported from Kolmogorov–Smirnov test (p= 9.56E-4, WT naïve L23: 15 cells, 6 mice L23; WT post: 28 cells, 7 mice). Bar graphs of averaged EPSCCRACM amplitudes ± SEM for WT pre- and post-training groups (inset). Significance was reported from the Mann–Whitney U test (p=1.059E-3). The average sCRACM maps across each grid were plotted below the corresponding cumulative. **(D)** Averaged EPSCCRACM amplitudes ± SEM by grid position in the vertical direction (perpendicular to the brain surface) from V1 L2/3 PCs in WT mice. Significance was reported from two-way ANOVA (genotype: F = 47.90; p = 1.737E-11). **(E)** Same as (B), but from FX pre- and post-training groups. Significance was reported from Kolmogorov–Smirnov test (p=1.658E-3, FX naïve L2/3: 23 cells, 6 mice; FX post-training L2/3: 30 cells, 8 mice). Significance was reported from the Mann–Whitney U test (p=0.0117). **(F)** Same as (C), but from FX pre- and post-training groups. In the vertical direction, significance was reported from two-way ANOVA (genotype: F = 14.64; p = 1.466E-4). **(G)** Illustration of V1 to LM L4 PC projections. **(H)** Same as (B), but from LM L4 PCs. Significance was reported from Kolmogorov–Smirnov test (p= 0.134, WT naïve L4: 6 cells, 4 mice; WT post-training L4: 15 cells, 6 mice). Significance was reported from the Mann–Whitney U test (p= 0.0233). **(I)** Same as (C), but from LM L4 PCs. In the vertical direction, significance was reported from two-way ANOVA (genotype: F = 10.46; p = 1.44E-3). **(J)** Same as D) but from LM L4 PCs. Significance was reported from Kolmogorov–Smirnov test (p= 0.517, FX naïve L4: 12 cells, 6 mice; FX post-training L4: 13 cells, 6 mice). Significance was reported from the Mann–Whitney U test (p= 1.0). **(K)** Same as (E), but from L4 PCs. In the vertical direction, significance was reported from two-way ANOVA (genotype: F = 0.0163; p = 0.8984). **(L)** Illustration of V1 to LM L5 PC projections. **(M)** Same as (B), but from LM L5 PCs. Significance was reported from Kolmogorov–Smirnov test (p= 0.0198, WT naïve L5: 12 cells, 6 mice; WT post-training L5: 21 cells, 6 mice). Significance was reported from the Mann–Whitney U test (p= 0.0175). **(N)** Same as (C), but from L5 PCs. In the vertical direction, significance was reported from two-way ANOVA (genotype: F = 32.32; p = 3.017E-8). **(O)** Same as D) but from LM L5 PCs. Significance was reported from Kolmogorov–Smirnov test (p= 0.283, FX naïve L5: 18 cells, 6 mice; FX post-training L5: 26 cells, 8 mice). Significance was reported from the Mann–Whitney U test (p= 0.237). **(P)** Same as (E), but from L5 PCs. In the vertical direction, significance was reported from two-way ANOVA (genotype: F = 0.0837; p = 0.773). Data were presented as mean ± SEM. Two-way ANOVA: #p<0.05, ##p<0.01, ###p<0.001, ####p<0.0001. Kolmogorov–Smirnov test and Mann–Whitney U test: *p<0.05, **p<0.01, ***p<0.001, ****p<0.0001

To measure inter-areal connectivity, we optically stimulated ChR2-positive (ChR2+) axon terminals from anterograde neurons in a 10 by 10 grid (0.67 mm by 0.67 mm) covering a square area centered at recipient neurons in LM L2/3 to L5 using a light patterned illuminator (Figure 4A). For each 5 ms light pulse at each pixel, excitatory post-synaptic currents (EPSCs) were recorded under a voltage clamp at −70 mV. All sCRACM recordings were conducted in the presence of 10 μM tetrodotoxin (TTX) and 50 μM 4-aminopyridine (4-AP), sodium, and potassium channel inhibitors, to block action potentials and poly-synaptic responses. The CRACM heatmap was plotted for EPSCsCRACM values for each grid of light stimulations. The average EPSCsCRACM values and cumulative distributions of average EPSCsCRACM (logarithmic scale) were also plotted for the averaged EPSCs from each neuron. CRACM heatmaps of L2/3 EPSC amplitudes in experienced WT and FX mice were hotter than those of naive groups. Kernel density plots of pixel averaged EPSC values, along with mean EPSC amplitudes, were analyzed using the Kolmogorov-Smirnov (KS) and Mann-Whitney U tests, respectively, and demonstrated a significant increase post- vs pre-training for both WT and FX mice (Figure 4C, E). However, the increase in the average EPSCsCRACM in FX mice was smaller than in WT mice (Figure 4C, E). The profile of the averaged EPSCsCRACM amplitude by grid position was greatly elevated after the visual experience in WT mice (Figure 4D). This elevation was decreased in FX mice, but it was still significant compared to naïve FX mice (Figure 4F). We detected no difference between the naive and experienced groups in either WT or FX in V1 layer 4 (L4) (Figure 4H-J). The post-training CRACM heatmaps of the layer 5 WT and FX mice looked hotter than their corresponding naive groups. Nevertheless, only WT groups reached statistical significance (Figure 4L, M). We found no significant difference between WT and FX within the naïve groups, but the average EPSCsCRACM was higher in WT than in FX after the visual experience. The increase in the average CRACM EPSCs in FX mice was lower than that of WT mice (Figure 4B, C). The average EPSCsCRACM post-training was higher in WT compared to FX mice (Figure 4E). These results suggest that the visual experience-dependent synaptic plasticity deficits in FX are laminar layer-specific. We did not detect any difference between the naive and post groups in V1 L4 of either WT or FX mice (Figure 4G-J). The averaged EPSCsCRACM amplitude profile by grid position increased after visual experience in WT mice but not FX mice (Figure 4I, K). The CRACM heatmap of L5 WT and FX mice post-training looked hotter than their corresponding naive groups, but the statistical results only supported the difference in WT mice (Figure 4M, O). The averaged EPSCCRACM amplitude by grid position for L5 PCs was much higher in experienced WT mice compared to the naïve group, but there was no such difference within the FX groups (Figure 4N, P). We found no significant difference between naïve WT and naïve FX mice. However, the average EPSCsCRACM and averaged EPSCCRACM amplitude by grid position was higher in WT mice compared to FX mice after visual experience in L2/3 and L5 PCs, but not L4 PCs (Figure S5). The sCRACM results demonstrated that visual experience strengthened the synaptic connectivity in the FFV1->LM pathway of WT mice, and this plasticity was substantially impaired in FX mice. These results validate our directed information functional connectivity analysis and suggest that the impaired inter-areal synaptic connectivity might underlie the attenuated familiarity-induced theta frequency oscillations in LM of experienced FX mice.

### Attenuated synaptic connectivity in FB LM→V1 pathway from LM to V1 following visual experience in FX mice

To investigate the synaptic connectivity of the FB LM to V1 pathway, we proceeded with the same visual experience and sCRACM setup, but instead of V1, we injected AAV1-ChR2-Venus into LM. Three weeks later, we performed visual stimulation and sCRACM experiments (Figure 5A, F, K). The same analysis was conducted for the data from the FB pathway. The average EPSCsCRACM from LM to V1 L2/3 PCs showed a trend of decrease after visual experience but did not reach statistical significance (Figure 5B, top insert). No difference was detected in the cumulative distributions of average EPSCsCRACM from LM to V1 L2/3 PCs (Figure 5B, top). The averaged EPSCCRACM amplitude by grid position was lower in experienced mice (Figure 5C), which agreed with the EPSCsCRACM heatmap (Figure 5B, bottom). The FX mice displayed a similar trend of decrease in these three different analysis approaches (Figure 5D top, E), but the difference between the heatmap was not as large as in WT groups (Figure 5D bottom). No shift was observed in the averaged EPSCCRACM amplitude by grid position (Figure 5E top), but we found it shifted away from LM or the peak was wider (Figure 5E bottom). The sCRACM analysis for LM to V1 L4 PCs showed no difference between naïve and post-experience groups in all statistical analyses for both WT and FX mice (Figure 5G-J). The heatmaps also appeared similar among the four groups (Figure 5G, I bottom). Visual training-induced rise in averaged EPSCCRACM amplitude in V1 L5 PCs was revealed by all analysis methods, including the heatmaps in WT mice (Figure 5L, M). The heatmap was hotter in V1 L5 PCs in experienced groups (Figure 5N top). Still, the cumulative distributions of average EPSCsCRACM and averaged EPSCsCRACM amplitude showed no significant difference (Figure 5N top). We found an increase in averaged EPSCsCRACM amplitude by grid position, but no shift in the profile was notable (Figure 5O). No statistical difference was detectable when comparing genotypes in either naïve or experienced groups (Figure S6). Overall, the connectivity strength was lower in the FB pathway suggesting that visual experience elicited synaptic plasticity, which was deficient in FX mice, although the differences were milder than in the FF V1→LM pathway. Interestingly, our results suggest that visual experience might reduce the receptive field size in L2/3 PCs of the FB LM→V1 pathway in FX mice, which means visual experience might rectify some atypical circuit connectivity in FX mice. The visual experience-induced synaptic plasticity of the inter-areal projections formed onto L5 PCs was most severely impaired in FX mice in both FF V1→LM and FB LM→V1 pathways compared to the projections to the superficial and middle laminar layers.

**Figure 5.**
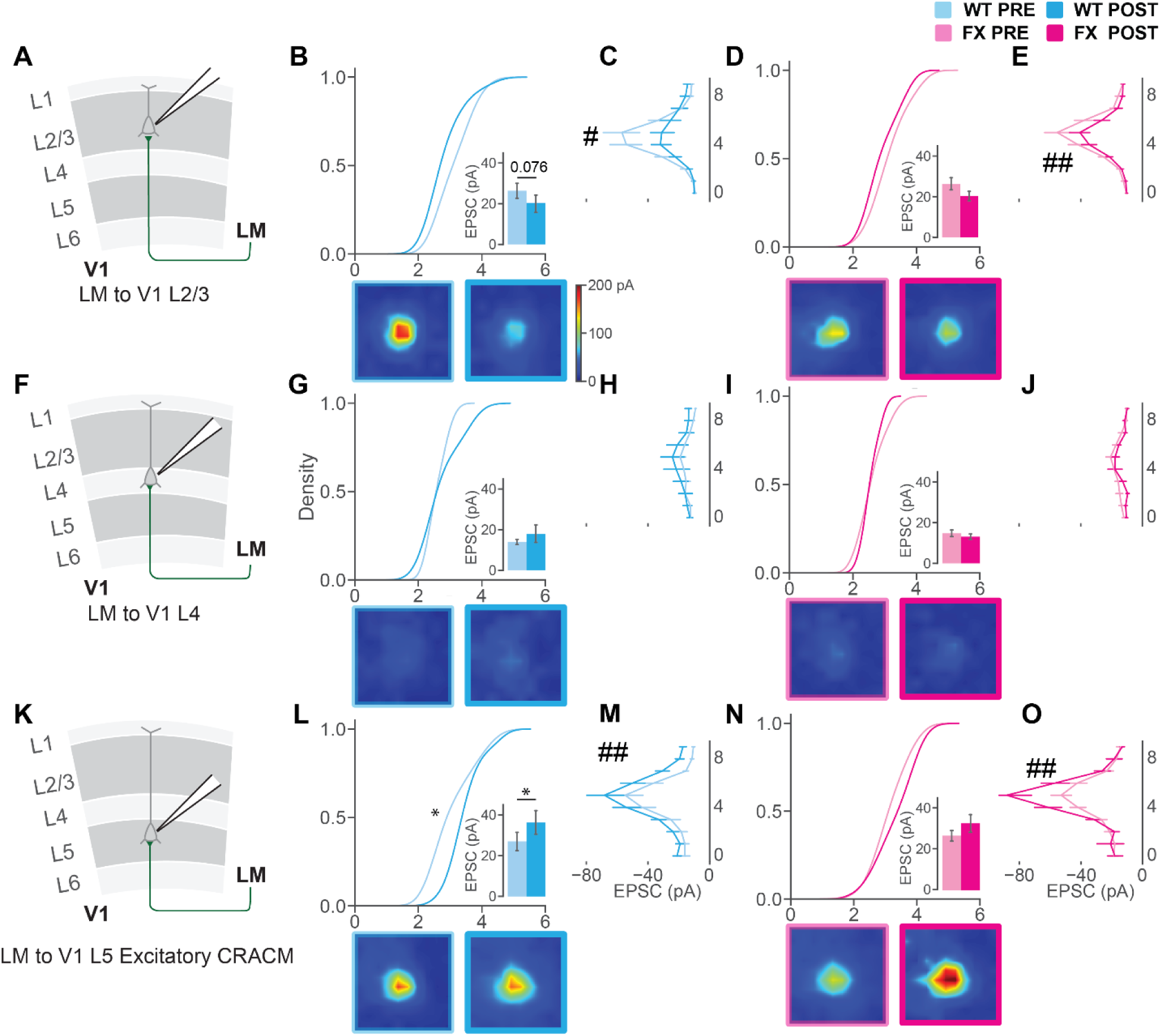
FB sCRACM maps of input from LM ChR2 positive neurons to PCs in V1. **(A)** Illustration of LM to V1 L23 PC projections. **(B)** The cumulative distributions (top) of the averaged EPSCCRACM amplitudes plotted in ln scale in sCRACM map recorded from LM L2/3 PCs. Significance was reported from the Kolmogorov– Smirnov test (p= 0.214, WT naïve L2/3: 19 cells, 6 mice; WT post-training L2/3: 17 cells, 6 mice). Bar graphs of averaged EPSCCRACM amplitudes ± SEM for WT pre and post-training groups (inset). Significance was reported from the Mann–Whitney U test (p= 0.0760). The average sCRACM maps across each grid were plotted below the corresponding cumulative. **(C)** Averaged EPSCCRACM amplitudes ± SEM by grid position in the vertical direction (perpendicular to the brain surface) from V1 L2/3 PCs in WT mice. Significance was reported from two-way ANOVA (genotype: F = 6.054; p = 0.0144 **(D)** Same as (B), but from FX pre- and post-training groups. Significance was reported from Kolmogorov–Smirnov test (p= 0.232, FX naïve L2/3: 37 cells, 16 mice; FX post L2/3: 26 cells, 9 mice). Significance was reported from the Mann–Whitney U test (p= 0.130). **(E)** Same as (C), but from FX pre and post-training groups. In the vertical direction, significance was reported from two-way ANOVA (genotype: F = 6.925; p = 8.713E-03). **(F)** Illustration of V1 to LM L4 PC projections. **(G)** Same as (B), but from LM L4 PCs. Significance was reported from Kolmogorov–Smirnov test (p= 0.754, WT naïve L4: 12 cells, 6 mice; WT post-training L4: 7 cells, 5 mice). Significance was reported from the Mann–Whitney U test (p= 0.967). **(H)** Same as (C), but from LM L4 PCs. In the vertical direction, significance was reported from two-way ANOVA (genotype: F = 3.77; p = 0.0540). **(I)** Same as D) but from LM L4 PCs. Significance was reported from Kolmogorov–Smirnov test (p= 0.910, FX naïve L4: 21 cells, 10 mice; FX post-training L4: 7 cells, 6 mice). Significance was reported from the Mann–Whitney U test (p= 0.836). **(J)** Same as (E), but from L4 PCs. In the vertical direction, significance was reported from two-way ANOVA (genotype: F = 2.469; p = 0.117). **(K)** Illustration of V1 to LM L5 PC projections. **(L)** Same as (B), but from LM L5 PCs. Significance was reported from Kolmogorov–Smirnov test (p= 0.0113, WT naïve L5: 21 cells, 8 mice; WT post-training L5: 17 cells, 7 mice). Significance was reported from the Mann–Whitney U test (p= 0.0492). **(M)** Same as (C), but from L5 PCs. In the vertical direction, significance was reported from two-way ANOVA (genotype: F = 10.95; p = 1.031E-03). **(N)** Same as D) but from LM L5 PCs. Significance was reported from Kolmogorov–Smirnov test (p= 0.209, FX naïve L5: 35 cells, 10 mice; FX post L5: 20 cells, 9 mice). Significance was reported from the Mann–Whitney U test (p= 0.231). **(O)** Same as (E), but from L5 PCs. In the vertical direction, significance was reported from two-way ANOVA (genotype: F = 8.559; p = 3.586E-03).

### Altered E/I balance of FF V1→LM and FB LM→V1 inputs to PCs in L2/3 and 5 in FX mice

We examined the circuit in a subcellular or mono-synaptic manner by applying TTX and 4AP, which abolished mixed synaptic inputs and di-synaptic involvement of inhibitory interneurons. However, *in vivo,* cortical network activity and modulation of neuronal responsiveness in a behaviorally relevant manner are achieved through a dynamic balance of excitation and inhibition ^40^. Many previous studies indicate that E/I balance is impaired in multiple models of ASDs ^41–45^. To measure excitatory and inhibitory synaptic balance (E/I balance) in PCs of FF V1→LM and FB LM→V1 pathways, we utilized CRACM as previously described ^46^. The glutamatergic synaptic inputs to PCs were recorded the same way as the sCRACM experiment, but without TTX and 4-AP, while the GABAergic synaptic inputs to PCs were recorded while holding the cells at the excitatory postsynaptic potential (EPSP) reversal potential. The compartmentalized E/I ratio recorded by CRACM, reflecting balanced synaptic strength in subcellular compartments, could shape subcellular circuitry and computation logic (e.g., soma vs. dendrites) ^46^. To investigate the E/I ratio at different subcellular locations for the FF V1→LM pathway in LM PCs in L2/3 and L5 (Figure S7A-D), we defined four grids surrounding the cell body of the recorded cell (indicated as a white triangle) as the ’soma’ area (Figure S9A, orange outline), and the outside grids as the ’dendrite’ area (Figure S9A, blue outline). The EPSC and IPSC traces of the whole neuron, soma, or dendrite were extracted to calculate the E/I ratio (Figure S9B, E/I ratio = peak of EPSC/peak of IPSC). The E/I ratio recorded in the LM L2/3 neurons in the FF V1->LM pathway did not alter pre- and post-experience in either WT or FX mice and showed no difference between WT and FX groups in neurons or divided compartments (soma or dendrite area, Figure S9C-E). The two compartments displayed similar E/I dynamics in the FF V1->LM pathway (Figure S9C-E). The average EPSCCRACM and IPSCCRACM and the cumulative distributions of averaged EPSCCRACM and IPSCCRACM showed no statistical difference after visual experience in either WT or FX mice (Figure S7E-H), which explained the consistency of the E/I ratio in V1 to LM L2/3 PCs pathway. The E/I ratio from the LM L5 neurons in the FF V1->LM pathway was lower in FX mice compared to WT mice before visual experience, but the ratio became comparable in post-experienced animals (Figure S9C-E). The visual experience moved the E/I balance toward excitation for FX mice but did not move it for WT mice (Figure S9C- D). The average EPSCCRACM and IPSCCRACM and the cumulative distributions of averaged EPSCCRACM and IPSCCRACM in L5 were both increased in WT mice, but only EPSCCRACM was increased in FX mice (Figure S7I-L). The soma and dendrite areas showed distinct E/I dynamics in the FB LM->V1 pathway, but L2/3 and L5 neurons displayed similar E/I dynamics induced by visual experience and for different genotypes. We found the E/I ratio was higher in naïve FX mice than in WT mice for the dendrite area and the whole neuron region, but no difference was detectable in the soma area (Figure S9F-H). In the opposite of the FF V1->LM pathway, the E/I balance moved toward excitation for WT mice after visual experience but not FX mice (Figure S9F-H). The average EPSCCRACM increased after the visual experience in both L2/3 and L5 neurons in WT mice, but IPSCCRACM showed no difference (Figure S8E, G, I, K), which was why we found that the E/I ratio increased in WT mice. EPSCCRACM and IPSCCRACM rose after visual experience in FX mice in L2/3 but not in L5 (Figure S8F, H, J, L). The results of circuit mapping without blocking action potentials were different from the sCRACM results because they included the feedforward excitation and inhibition of the local cortical circuits (Figure S7A-D, Figure S8A-D). Overall, the FF V1->LM and FB LM->V1 pathways showed distinct E/I dynamics in the superficial and deep layers and differences between distinct cellular compartments only in the FB LM->V1 pathway. Interestingly, the visual experience shifted the E/I balance and made it comparable between WT and FX mice in the FF V1->LM and FB LM->V1 pathways.

### Impaired STP in FF V1→LM and FB LM→V1 pathway in FX mice

To measure presynaptic STP, we used optogenetic paired-pulse stimulation of the synapses between V1 and LM with TTX and 4-AP in bath ^47–49^ with full-field light stimulation delivered through the 10x objective lens. We calculated the paired-pulse ratios (PPR=EPSC2/EPSC1) for a range of inter-pulse intervals (IPI) from 25 ms to 2000 ms. Short IPI stimulations induced depression in all animals, but this depression was decreased after visual experience in both WT and FX mice (IPI=50 ms, Figure 6A). The paired-pulse depression was dominant in both FF and FB corticocortical synapses between V1 and LM, consistent with other reports in the frontal cortex^50^. The PPRs increased after visual experience in L2/3 neurons from the FFV1->LM pathway of WT mice, especially for IPI’s: 50, 100, 150, 200, and 2000 ms (Figure 6B). There was no significant difference between naïve and experienced FX mice (Figure 6C). The PPR was larger in visually experienced FX mice only for IPI=50 ms; the PPR in the rest of the IPIs was either lower or similar compared to naïve mice (Figure 6C). When comparing the genotypes for the L2/3 neurons of the FFV1->LM pathway, naïve FX mice showed larger PPR than the corresponding WT mice, which was reversed after the visual experience (Figure S10A, B). In L5 neurons of the FFV1->LM pathway, the post hoc test indicated higher PPRs when IPI =25 and 50 ms in experienced mice for both WT and FX (Figure 6D, E). However, the ANOVA test only showed significant differences between pre- and post- experienced groups in WT mice but not FX mice (Figure S10D, E). No difference was detected between WT and FX in naïve groups (Figure S10C). Like L2/3 neurons, we saw that PPRs were generally larger in WT than FX in post-visual experience groups (Figure S10D).

**Figure 6.**
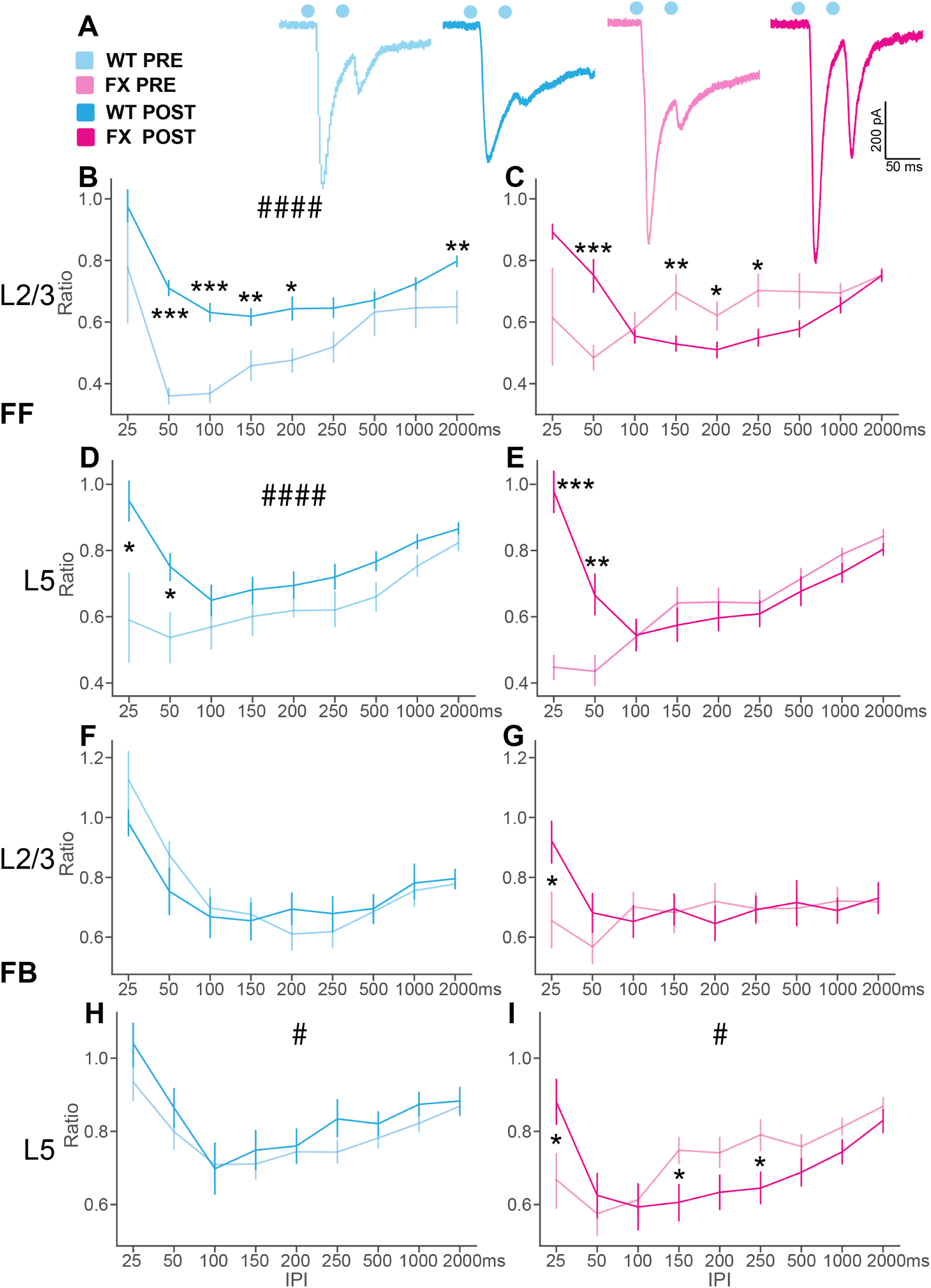
Paired pulse facilitation of FF V1 to LM and FB LM to V1 L2/3 and L5 PCs projections comparing pre and post-perceptual experience. **(A)** Example EPSCs elicited by paired-pulse paradigm (50 ms IPI) in four groups of animals. Blue dots highlighted the time when optical stimulation was delivered. **(B)** Average PPRs of naïve and post-training WT mice of corresponding inter-pulse intervals in FFV1->LM L2/3 pathway (WT pre: N=65 cells, WT post: N=195 cells). Significance was reported from two-way ANOVA (perceptual experience: F = 2.757; p = 2.104E-3) and Tukey’s HSD tests (25ms, p= 0.2011; 50 ms, p= 0.001; 100 ms p= 0.001; 150 ms, p= 0.0084; 200 ms, p=0.0187; 250 ms, p= 0.0551, 500 ms, p= 0.5569; 1000 ms, p= 0.1554; 2000 ms, p= 0.0018). **(C)** Same as (B), but from FX mice (FX pre: N=158 cells, FX post: N=169 cells). Significance was reported from two-way ANOVA (perceptual experience: F = 2.757; p = 0.0978) and Tukey’s HSD tests (25 ms, p= 0.1984; 50 ms, p= 0.001; 100 ms p= 0.0262; 150 ms, p=0.1679; 200 ms, p= 0.0432; 250 ms, p= 0.0132, 500 ms, p= 0.0551; 1000 ms, p= 0.3333; 2000 ms, p= 0.9). **(D)** Average PPRs of naïve and post-training WT mice of corresponding inter-pulse intervals in FFV1->LM L5 pathway (WT pre: N=97 cells, WT post: N=209 cells). Significance was reported from two-way ANOVA (perceptual experience: F = 25.35; p = 8.42E-7) and Tukey’s HSD tests (25ms, p= 0.0108; 50 ms, p= 0.0124; 100 ms, p= 0.348; 150 ms, p= 0.2762; 200 ms, p= 0.3041; 250 ms, p= 0.1317, 500 ms, p= 0.0559; 1000 ms, p= 0.066; 2000 ms, p= 0.1866). **(E)** Same as (D), but from FX mice (FX pre: N=148 cells, FX post: N=183 cells). Significance was reported from two-way ANOVA (perceptual experience: F = 2.60; p = 0.108) and Tukey’s HSD tests (25ms, p= 0.001; 50 ms, p= 0.0073; 100 ms, p= 0.9; 150 ms, p= 0.3509; 200 ms, p= 0.4762; 250 ms, p= 0.5895, 500 ms, p= 0.4863; 1000 ms, p= 0.1385; 2000 ms, p= 0.1349). **(F)** Average PPRs of naïve and post-training WT mice of corresponding inter-pulse intervals in FBLM->V1 L2/3 pathway (WT pre: N=126 cells, WT post: N=119 cells). Significance was reported from two-way ANOVA (perceptual experience: F = 7.69E-3; p = 0.930) and Tukey’s HSD tests (25ms, p= 0.206; 50 ms, p= 0.234; 100 ms p= 0.774; 150 ms, p= 0.827; 200 ms, p= 0.284; 250 ms, p= 0.447, 500 ms, p= 0.880; 1000 ms, p= 0.759; 2000 ms, p= 0.757). **(G)** Same as (F), but from FX mice (FX pre: N=221 cells, FX post: N=214 cells). Significance was reported from two-way ANOVA (perceptual experience: F = 2.757; p = 0.0978) and Tukey’s HSD tests (25ms, p= 0.050; 50 ms, p= 0.199; 100 ms p= 0.508; 150 ms, p= 0.877; 200 ms, p= 0.400; 250 ms, p= 0.9, 500 ms, p= 0.849; 1000 ms, p= 0.643; 2000 ms, p= 0.848). **(H)** Average PPRs of naïve and post-training WT mice of corresponding inter-pulse intervals in FBLM->V1 L5 pathway (WT pre: N=248 cells, WT post: N=161 cells). Significance was reported from two-way ANOVA (perceptual experience: F = 4.76; p = 0.0297) and Tukey’s HSD tests (25ms, p= 0.167; 50 ms, p= 0.367; 100 ms p= 0.886; 150 ms, p= 0.582; 200 ms, p= 0.782; 250 ms, p= 0.113, 500 ms, p= 0.403; 1000 ms, p= 0.200; 2000 ms, p= 0.759). **(I)** Same as (H), but from FX mice (FX pre: N=267 cells, FX post: N=214 cells). Significance was reported from two-way ANOVA (perceptual experience: F = 4.04; p = 0.045) and Tukey’s HSD tests (25ms, p= 0.048; 50 ms, p= 0.587; 100 ms p= 0.800; 150 ms, p= 0.029; 200 ms, p= 0.0960; 250 ms, p= 0.0211, 500 ms, p= 0.160; 1000 ms, p= 0.102; 2000 ms, p= 0.338). Data were presented as mean ± SEM. Two-way ANOVA: #p<0.05, ##p<0.01, ###p<0.001, ####p<0. 0001.Tukey HSD’ test: *p<0.05, **p<0.01, ***p<0.001, ****p<0.0001

Consistent with the sCRACM results for the FB LM->V1 pathway, the PPR recorded in L2/3 did not differ much between naïve and post-experienced mice in both WT and FX (Figure 6F, G). Visual experience enhanced the PPRs recorded in L5 for WT mice but reduced it for FX mice when IPI=150 ms and 250 ms (Figure 6H, I). The post hoc test showed an increase in PPR when IPI=25 ms for neurons recorded in L2/3 and L5 (Figure 6G, I). When comparing genotypes in the FB LM->V1 pathway, the PPR was larger when IPI was low in naïve WT mice than in post-visual experienced mice in neurons recorded in both L2/3 and L5 revealed by post hoc test (Figure S10E, G). However, the ANOVA test only indicated a difference in neurons recorded in L5 (Figure S10G). In post-visual experienced groups, the PPRs recorded in L2/3 of the FB LM->V1 pathway were comparable between WT and FX, but it was larger in WT mice in L5 (Figure S10H). Overall, the visual experience led to an increase in PPRs, especially for the FF connections from V1 to LM in WT mice, and to a lesser extent in FX mice.

**Experience-dependent structural plasticity of dendritic spines in L5 PCs was impaired in FX mice.**

Previous studies reported abnormal dendritic spine phenotypes in the superficial cortical L2/3 neurons, including increased spine density and relative immaturity of spines in adult FX mice ^11,51–54^. However, other research reported no changes in the dendritic spine morphology ^11,55–57^. This discrepancy in the reports may result from the limitation of the confocal and two-photon imaging techniques with relatively low resolution of the dendritic spines. Single-molecule localization microscopy (SMLM) ^58,59^ has overcome the diffraction barrier and provided unprecedented opportunities to observe cellular functions, interactions, and dynamics at the nanoscale level ^60^. Specifically, SMLM (also known as PALM/STORM), as well as its three-dimensional (3D) counterpart ^61,62^, utilizes photo-switchable or convertible dyes to allow the detection and localization of isolated molecules with a precision as low as 5 nm in 3D. We have recently developed a new SMLM method for thick tissues providing an opportunity to perform super-resolution imaging of synaptic ultrastructures in brain slices with preserved intact neural circuits.

To image the dendritic spines of the L5 PCs in naïve and experienced WT and FX mice, we performed an in vivo visual stimulation paradigm as described earlier using a double transgenic cross of Fmr1 KO line and Thy1-ChR2-EYFP ^63^. After sacrificing and processing the mouse brains for SMLM microscopy, we performed blind imaging of the dendritic spines of L5 PCs in all four different conditions: naïve Thy1-ChR2-EYFP, naïve Fmr1 KO x Thy1-ChR2-EYFP, experienced Thy1-ChR2-EYFP, experienced Fmr1 KO x Thy1-ChR2-YFP (Figure 7A, B). To measure the size of dendritic spines of apical dendrites from L5 PCs, we developed an approach to clearly resolve and quantify the contour of a cross-section of the spine, neck, and head using SMLM (Figure 7C), allowing precise fitting and estimating the size of the dendritic spines (Figure S11). As previously reported, the spine density was higher in FX mice than in WT mice in naïve groups (Figure 7D). We found that the visual experience greatly affected the plasticity of the protrusion density in WT mice but did not affect the density in FX mice (Figure 7D). The dendritic spines were longer in naïve FX mice than the naïve WT mice, but no difference was detected between genotypes in experienced groups (Figure 7E). The visual experience did not alter the length of the protrusions for either WT or FX mice in the visual cortex (Figure 7E). The spine volumes remained stable in WT and FX mice following the visual experience (Figure 7F). The size of the spine neck (minimal spine area) was larger in naïve WT mice compared to FX mice (Figure 7G). When comparing the head size (maximal spine area), we found that visual familiarity upregulated the spine head size, and the post hoc test indicated statistical significance only in WT mice (Figure 7H). Overall, the spines in L5 PCs of the visual cortex of FX mice displayed relatively immature features (higher density, longer, thinner neck) and were insensitive to modulations by sensory experience.

**Figure 7.**
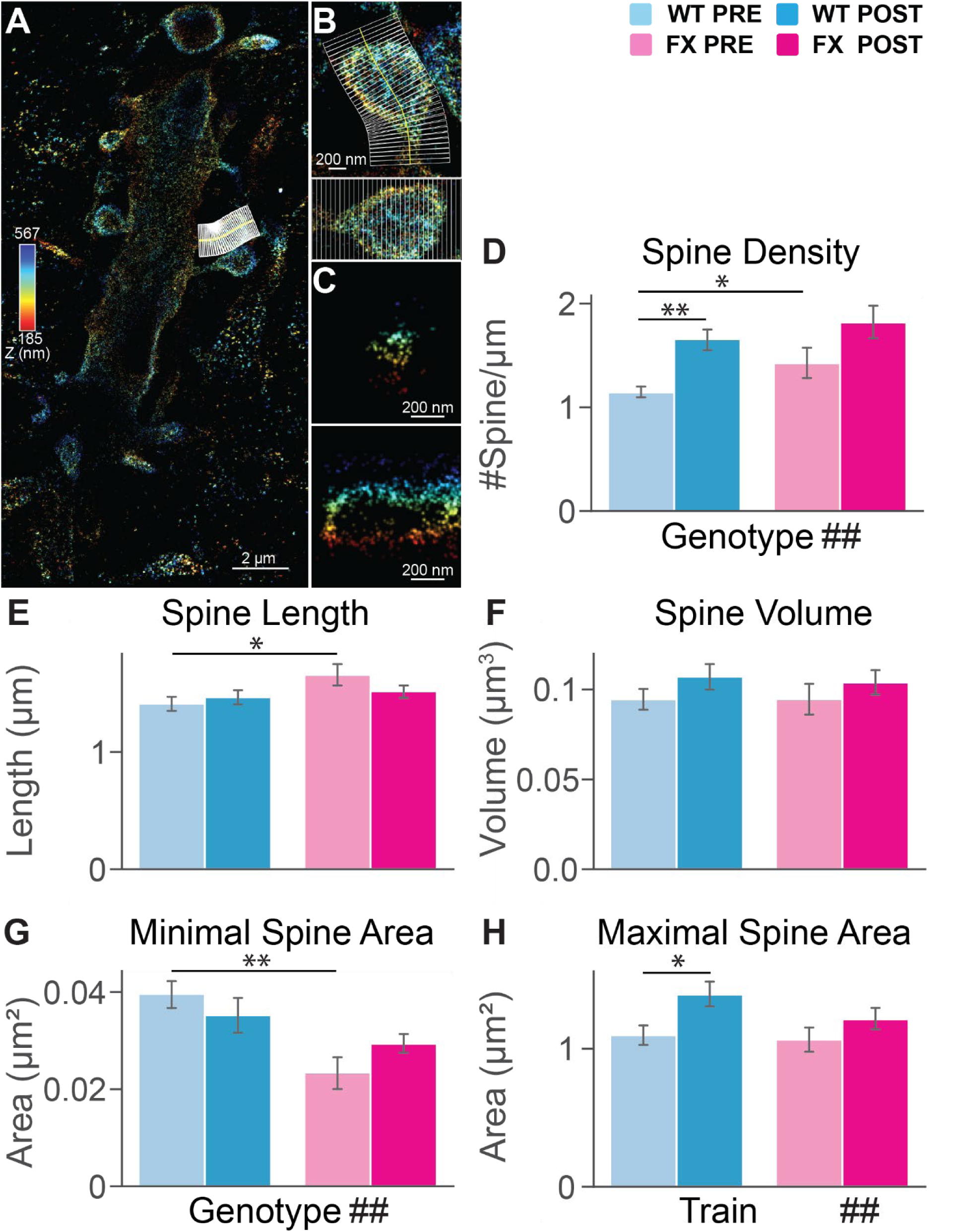
Spine morphology of V1 L5 PC dendrites. **(A)** An example of a 4Pi image with a dendritic spine, pseudo-colored based on localized axial position, selected for measurement. **(B)** The selected dendritic spine in (A) (Top). The straightened dendritic spine (bottom). **(C)** Axial cross-sections of the selected dendritic spines in (A) (top: neck area; bottom: head area). **(D)** Bar graphs of average spine length from dendrites of V1 L5 PCs in WT and FX mice (WT pre: N=81, WT post: N=95, FX pre: N=41, FX post: N=108 spines). Significance was reported from two-way ANOVA followed by Tukey’s HSD tests (two-way ANOVA: genotype: F = 3.13; p = 0.078. Perceptual experience: F = 0.126; p = 0.72. Tukey’s post hoc: WT pre versus FX pre: p = 0.033; WT post versus FX post: p = 0.53; WT pre versus WT post: p = 0.55; FX pre versus FX post: p = 0.19). **(E)** Bar graphs of average spine volume from dendrites of V1 L5 PCs in WT and FX mice. Significance was reported from two-way ANOVA followed by Tukey’s HSD tests (two-way ANOVA: genotype: F = 1.5E-5; p = 1.0. Perceptual experience: F = 2.27; p = 0.13. Tukey’s post hoc: WT pre versus FX pre: p = 0.9; WT post versus FX post: p = 0.75; WT pre versus WT post: p = 0.18; FX pre versus FX post: p = 0.45). **(F)** Bar graphs of average minimal spine area from dendrites of V1 L5 PCs in WT and FX mice. Significance was reported from two-way ANOVA followed by Tukey’s HSD tests (two-way ANOVA: genotype: F = 10.20; p = 1.55E-3. Perceptual experience: F = 5.34E-3; p = 0.94. Tukey’s post hoc: WT pre versus FX pre: p = 1E-3; WT post versus FX post: p = 0.15; WT pre versus WT post: p = 0.35; FX pre versus FX post: p = 0.11). **(G)** Bar graphs of average maximal spine area from dendrites of V1 L5 PCs in WT and FX mice. Significance was reported from two-way ANOVA followed by Tukey’s HSD tests (two-way ANOVA: genotype: F = 0.95; p = 0.33. Perceptual experience: F = 5.34E-3; p = 9.30E-3. Tukey’s post hoc: WT pre versus FX pre: p = 0.79; WT post versus FX post: p = 0.14; WT pre versus WT post: p = 0.013; FX pre versus FX post: p = 0.29). **(H)** Bar graphs of average maximal spine area from dendrites of V1 L5 PCs in WT and FX mice (WT pre: N=17 dendrites, WT post: N=10 dendrites, FX pre: N=11 dendrites, FX post: N=11 dendrites). Significance was reported from two-way ANOVA followed by Tukey’s HSD tests (two-way ANOVA: genotype: F = 8.58; p = 5.33E-3. Perceptual experience: F = 2.61; p = 0.11. Tukey’s post hoc: WT pre versus FX pre: p = 0.048; WT post versus FX post: p = 0.44; WT pre versus WT post: p = 1E-3; FX pre versus FX post: p = 0.091). Data were presented as mean ± SEM. Two-way ANOVA: #p<0.05, ##p<0.01, ###p<0.001, ####p<0.0001. Tukey HSD’ test: *p<0.05, **p<0.01, ***p<0.001, ****p<0.0001

## Discussion

### Impaired visual familiarity-dependent theta oscillations and synchrony in V1 and LM of FX mice

Theta frequency oscillations in cortical regions have been recognized as associated with visual familiarity ^8,33,36,64^. Previously, we demonstrated that these oscillations within V1 were impaired in FX mice, characterized by a shifted frequency, reduced power, and shorter duration ^9^. These oscillations might represent a communication mechanism between V1 and HVAs, including top-down modulation of V1, as has been shown in previous studies ^35,65,66^. Visual familiarity-induced 4–8 Hz oscillations might represent the mechanism by which V1 and HVAs synchronize across different brain areas, facilitating the processing of familiar content ^35^. Consistent with our previous studies, we discovered oscillatory activity in V1 and LM that became more phase synchronized post visual experience in a stimulus-selective feature-dependent manner in WT mice^35^. In FX mice, the visual experience evoked 4-8 Hz theta frequency oscillations in V1 and LM were both attenuated.

LFP analysis showed a loss in power of the 4-8 Hz oscillations for both V1 and LM of FX mice compared to WT mice. Despite choosing a visual stimulus that was shown to elicit the strongest response in LM, phase synchronization of LFP peaks was only seen in WT mice, indicating a loss of inter-areal connectivity between V1 and LM of FX mice. Impairments in the oscillatory activity of FX mice were also represented in the unit population firing rates of V1 and LM, with decreased firing rates at the oscillation cycles following the stimulus presentation time window in FX mice. Directed information analysis revealed significant differences in the functional connectivity of V1 and LM post-visual experience between WT and FX mice. Most notably, there was an overall increase in FF input from V1 to LM in WT mice post-training but not in FX mice. Further, there was a slight increase in the feedback input from LM to V1 post-visual experience in WT mice, while this feedback input was decreased in FX mice. These differences in functional connectivity indicate deficits in communication between V1 and HVAs of FX mice.

### Impaired V1-LM inter-areal synaptic connectivity after visual experience in FX mice

Our circuit mapping studies demonstrated uneven deficits in synaptic connectivity across FF _V1→LM_ and FB LM→V1 pathways induced by visual experience in FX mice and validated our functional connectivity results. Specifically, we observed a significant strengthening of connections from V1 to the superficial, middle, and deep layer neurons in LM after visual experience in WT mice. In contrast, in FX mice, this enhancement was confined to the superficial layers and to a lesser extent compared to WT. However, the alterations in connectivity strength triggered by visual experience in the FB pathway were more subtle in WT mice, while in FX mice, these deficits were mild. Additionally, we found that visual experience elicited synaptic plasticity in a layer-specific manner for both FF V1→LM and FB LM→V1 pathways, with varying degrees of impairment across cortical layers in FX mice. These distinctions may be attributed to the laminar preferences for cortico-cortical connectivity due to the hierarchical organization of areas in the mouse visual cortex ^24,67^. In rodents, projections from V1 terminate from inner L1 to out L6, while FB LM→V1 projections from the extrastriate cortex to V1 typically avoid L4, preferentially targeting layers L1, L2/3, L5, and L6 ^24^. In previous work, we reported synaptic strengthening from L5 projections onto L2/3 and L4 after visual experience in V1, as evidenced by in vivo patch-clamp recordings combined with optogenetics, aligning with our FF V1→LM circuit mapping results ^36^. The uneven deficits observed were consistent with our findings that interareal reciprocal connectivity between V1 and LM exhibits distinct strength according to laminar specificity, providing direct evidence supporting previous studies that demonstrated distinct connections and response properties between V1 and HVAs ^22,68,69^. Interestingly, our results are also consistent with our previous work describing layer-specific alterations in visual responses to the oddball paradigm in FX mice, potentially providing insights into the circuit mechanisms underlying these impairments. ^70^.

The visual training paradigm specific for the ventral path elicited elevation in Fos-positive neuron number in WT mice in LM of the ventral pathway but not in AL, AM, or PM of the dorsal pathway because this specific visual stimulus maximally induced response in LM but not in the dorsal pathway^35^. The number of Fos-positive neurons in V1 was almost ten times that in LM (Figure 1), and the excitatory postsynaptic currents measured by CRACM for the FF V1→LM pathway were also significantly higher than those in the FB LM→V1 pathway (Figure 4 and Figure 5). The weaker FB LM→V1 projections we detected are consistent with findings by Burkhalter, A. et al.^68^. Previous studies revealed that FX mice displayed altered hippocampal synaptic plasticity, including enhanced augmentation, but not short-term facilitation, in hippocampus synapses, indicating abnormal STP during high-frequency stimulations^16,71^. We also found that the STP in synapses from distinct layers of FF V1→LM and FB LM→V1 pathways was not consistently altered in naïve FX mice (Figure S10). However, the experience-dependent elevation of PPRs was defective in FX mice, and PPRs were generally lower in FX mice after the visual experience, especially in synapses of the FF V1→LM pathway (Figure 6 and Figure S10). The visual experience-dependent STP changes in WT mice and the abnormal STP in FX mice aligned well with the synaptic strength increase in sCRACM results, which implies a pre-synaptic mechanism in visual familiarity-induced synapse plasticity in the FF V1→LM and FB LM→V1 pathways. There is a scarcity of studies focused on the inter-areal interaction and sensory binding in ASDs ^72^. Our results may provide valuable insights into the mechanisms of interareal interaction in the visual cortex that underlie visual learning deficits in individuals with autism addressing a significant gap in the existing research.

### FX and dendrite morphology in visual familiarity

Dendritic spines exhibit a strong potential for morphological plasticity, enabling neurons to modify synaptic connectivity, adaptively remodel neural circuits, and support learning and memory processes ^73^. Sensory experiences, even in adulthood, can drive the formation and elimination of synapses ^74^. In typical development, the dendritic spine becomes shorter, the heads of the dendritic spine become smaller, and the necks widen ^75,76^. The visual familiarity paradigm can also induce morphological plasticity, resembling the developmental trajectory of the dendritic spine. Cell-autonomous removal of Fmr1 postnatally in postsynaptic L2/3 or L5 neurons leads to a specific reduction in the strength of callosal synaptic function mediated by AMPA receptors (AMPARs), while functions mediated by NMDA receptors remain unaffected, suggesting the presence of immature synapses ^77^. However, comparisons between P14 and P37 mice using STED microscopy revealed only subtle differences between WT and FX mice ^76^. SMLM achieved much higher resolution than STED^78^ and allowed us to assess dendritic spine density and morphology in V1 L5 pyramidal cells (PCs) in three to four-month-old WT and FX mice. We observed significant differences in dendritic spine plasticity: FX mice exhibited higher spine density, longer spines, and narrower spine necks compared to WT. Visual experience increased spine density and enlarged spine heads in WT mice, but no changes were observed in FX mice. The enhanced synaptic connectivity in L5 PCs in WT mice could be partially explained by the more mature features of the dendritic spines we observed.

### E/I imbalance and presynaptic plasticity disruption may contribute to the attenuated oscillations in FX mice

We observed an E/I imbalance in the FF V1→LM and FB LM→V1 pathways of FX mice, suggesting distinct deficits in circuit mechanisms underpinning top-down and bottom-up visual information processing. Specifically, the FF pathway exhibited inhibition, while the FB LM→V1 pathway demonstrated hyperexcitability in FX mice. The hyperexcitation in naïve mice may be attributed to aberrant connectivity in LM and the overall rise of neural activities in the visual areas, including certain laminar layers in V1 and LM, revealed by c-fos level quantification (Figure 1 and Figure S1). Interestingly, visual experience led to an increase in the excitatory drive in the FB LM→V1 pathway in WT mice but did not alter the E/I ratio in the FF V1→LM pathway (Figure S9). Conversely, the same visual experience elicited opposite effects on the E/I balance in FX mice (Figure S9), indicating that visual experience-induced plasticity is compromised in the inner-laminar connectivity in V1 of FX mice ^9^. This impairment may potentially explain the lack of excitation in the FB LM→V1 pathway following visual experience in FX mice. Despite the imbalance in the E/I ratio of FX, visual training might hold promise for at least partially rectifying the atypical neural connectivity, particularly for the FF V1→LM and FB LM→V1 pathways we examined. This strategy holds promise and is supported by other research demonstrating that evoked activity in PV+ interneurons in FX mice via a chemogenetic strategy leads to a potential rescue effect in E/I balance and may also compensate for the circuit disruptions observed in FX^79^.

### The promise of visual training as a neural circuit-based therapy for ASD patients

The E/I balance and STP experiments suggest that visual training may alleviate at least some of the atypical synaptic and circuit anomalies in FX. Previous studies have demonstrated that environmental enrichment can facilitate behavioral and neuronal morphological recovery in FX mice^80,81^ and improvement in social performance and learning in ASD children ^82–85^. Individuals with ASD showed a preference for processing low spatial frequency content rather than high spatial frequency content in face categorization strategies ^86–88^ together with a deficit in visual temporal integration^89,90^. These studies and our research support the notion that the visual perception of ASD patients is atypical in terms of processing spatial and temporal frequency information of the visual content. Our findings suggest that repetitive training with the specific patterns of spatial and temporal frequencies could potentially be beneficial to FX mice and might also have therapeutic indications for ASD patients. Future research should explore a broader range of spatial and temporal frequency patterns in visual training to determine which configurations yield the most effective behavioral interventions ^35^. Such investigations could pave the way for innovative forms of visual therapy for individuals with ASD.

## Acknowledgement

We thank the Chubykin lab members for valuable comments. We thank Dr. Sotiris Masmanidis for providing silicon probes. This work was supported by the US National Institutes of Health (grants AA027301 and AA029985 to A.K., GM119785 to F.H., MH123401 to F.H. and A.A.C., MH116500 to A.A.C.). F.X. acknowledges the National Key R&D Program of China (2023YFC3402600) and the National Natural Science Foundation of China (62272041).

## Author Contributions

X.C. and A.A.C. conceptualized the study. X.C., S.N., H.C.G, Y.C., T.X., F.X., P.A.E. performed experiments and analyzed data. Y.Y.N. and C.J.Q. analyzed data. Y.X.C. and S.N. wrote the original draft. All authors reviewed and edited the manuscript. A.K., F.H. and A.A.C. supervised the experiments, data analysis and writing.

## Methods

### Experimental Model and Subject Details

All experimental procedures were approved by the Purdue Animal Care and Use Committee (PACUC, protocol number 140800111. Three-to four-month-old homozygous male and female Fmr1 KO (B6.129P2-Fmr1tm1Cgr/J, JAX Stock No. 003025), B6 (C57BL/6, JAX), and Thy1-ChR2-YFP (B6.Cg-Tg(Thy1-COP4/EYFP)18Gfng/J, JAX Stock No. 007612) were used for the experiment.

### Visual experience paradigm

Mice were habituated to the training apparatus before passive visual experience training, which followed two weeks post-surgery. During the 2-3 days of habituation, a monitor displaying a grey screen was placed 17cm in front head-fixed mice in a dark environment for an hour/day. For passive visual training, a filtered pink noise picture of 0.12 cycles/degree was displayed, with the illuminance intensity of each pixel altered in a sinusoidal function of time at 0.75 Hz to generate a movie. Filtered pink noise movie stimulus was displayed using PsychoPy. Mice were shown 200 trials of 200ms pink noise movie stimulus, with a 6-sec grey screen between trials, for 4-6 days. For *in vivo* recordings, mice were presented with 20 stimulus trials on recording day.

### iDISCO whole-brain immunostaining

The brains were first perfused and collected 90 minutes after the visual experience paradigm was conducted. They were then stored in a 4% PFA solution for 24 hours, followed by three washes with 1x PBS for 30 minutes each. The samples are processed following the protocol from the previous literature ^91–95^. The fixed samples underwent a series of washes in methanol of increasing concentrations: 20%, 40%, 60%, 80%, and 100%, each for one hour, with two repetitions at 100% methanol. Subsequently, the samples were incubated overnight in a 66% dichloromethane (DCM) solution and 33% methanol. The next day, the samples were washed twice with 100% methanol for one hour each. To bleach the brains, they were exposed overnight to a solution of 5% hydrogen peroxide in methanol at 4 °C overnight. The next day, the samples underwent a rehydration process using a reverse series of methanol concentrations ranging from 100% to 20% in double-distilled water (ddH2O), with additional one-hour washes in PBS. Next, the brains underwent two one-hour washes in PBS/0.2% Triton-X-100, followed by two days of permeabilization at 37°C in a solution composed of 0.3M glycine/20% dimethyl sulfoxide (DMSO)/1x PBS with 0.2% Triton-X-100. Subsequently, the samples were blocked for two days at 37°C in a solution of PBS containing 0.2% Triton-X-100, 10% DMSO, and 6% normal donkey serum. The brains were then incubated for seven days in a primary antibody solution (rabbit anti-cFos, 1:1000; Synaptic Systems #226-003) diluted in PBS containing 0.2% Tween-20, 5% DMSO, 3% normal donkey serum, and 10 μg/mL heparin. The samples were then washed in PBS containing 0.2% Tween-20 and 10 μg/mL heparin (PtWH) over 24 hours, with 5 changes of solution. The brains were incubated in a secondary antibody solution (donkey anti-rabbit Alexa Fluor 647, 1:500; Thermo Fisher Scientific #A-31573) diluted in PBS containing 0.2% Tween-20, 5% DMSO, 3% donkey serum, and 10 μg/mL heparin for seven days. Subsequently, the brains were washed with PtWHfor 24 hours as before, followed by dehydration using a decreasing series of methanol concentrations in ddH2O (20% to 100%), as previously described. The following day, the brains were incubated in a solution of 66% DCM/ 33% methanol for three hours and then two times in 100% DCM for 15 minutes each. After the DCM incubation, the brains were transferred to a clean tube containing dibenzyl ether until they were ready for imaging.

### Light sheet microscope image acquisition

The right hemispheres of the iDISCO+ processed samples were positioned for sagittal imaging (with the right lateral side facing up) using an UltraMicroscope II light-sheet microscope (Miltenyi Biotec, Bielefeld, Germany) equipped with a scientific complementary metal–oxide–semiconductor camera (Andor Neo), a 2×/0.5 objective lens (MVPLAPO 2×), and a dipping cap with a working distance of 6mm. The image capture was facilitated by InspectorPro software version 7.1.4. The microscope was equipped with an NKT Photonics SuperK EXTREME EXW-12 white light laser with three fixed light-sheet-generating lenses on each side. Scans were made at 0.8× magnification (1.6× effective magnification) with a light-sheet numerical aperture of 0.148. Excitation filters of 480/30 and 560/40 were used. Emission filters of 525/50 and 595/40 were used. The samples were scanned with a step size of 3 μm for the 480 and 560 channels (1 acquisition per plane with 560 ms exposure. Laser power was set at 55% for 480 channels and 10% for 560 channels).

### Identification of Activated Brain Regions

Images that were acquired from the light-sheet microscope were analyzed from the end of the olfactory bulbs (the olfactory bulbs were not included in the analysis) to the beginning of the hindbrain and cerebellum. Counts of Fos-positive nuclei from each sample were identified for each brain region using ClearMap2 ^94,96^. ClearMap uses autofluorescence acquired in the 488 channel to align the brain to the Allen Mouse Brain Atlas and then registers Fos counts to regions annotated by the atlas. Fos counts of midbrain reticular nucleus (MRN), primary visual area (VISP), and lateral visual area (VISl) were quantified.

### Head plate installation and virus injection

For surgeries of virus injection, head cap, or head plate implantation, mice were anesthetized by inhaling 1.5-2.0% isoflurane anesthesia and fixed in a stereotaxic apparatus. The location of V1 (3.0 mm lateral of midline, 0.8 mm anterior of transverse sinus [TS) or higher visual area LM (4.0 mm lateral of midline, 1.4 mm anterior of TS) were stereotaxically identified and marked on the skull by a marker pen. For injections, the skull on the marked sites was thinned by a dental drill to the extent that it can be penetrated by an injection glass pipette, tip diameter 20 μm. Intracerebral injections were conducted by Nanoject III Injector. The glass pipette was inserted stereotaxically 0.4 and 0.7 mm underneath the pial surface. 50 nL pACAGW-ChR2-Venus-AAV1 (Addgene cat#20071, titer ≥ 1×10¹³ vg/mL) was released to express ChR2 throughout the cortical layers and their outgoing axons for a long-range CRACM experiment measuring reciprocal connectivity in the visual cortex (FBLM→V1, FFV1 →LM). A customized head plate was placed on top of the skull. The skull and the head plate were covered with Metabond (Parkell S380). To control the ChR2 expression level, we injected an adequate amount of the virus, and we waited for virus expressions to plateau two to three weeks after infections before habituation. For *in vivo* recordings, an additional reference pin was attached to the skull and the head plate.

### Data acquisition and analysis of extracellular recordings

*In vivo,* simultaneous extracellular recordings were taken using two 64-channel silicon probes. Craniotomies were performed on the mouse over V1 and LM regions on the stereotaxic surgery apparatus under isoflurane gas. The mouse was then head-fixed, and the two probes were placed over the craniotomies^97^ and inserted at V1 and LM using micromanipulators (NewScale). Before insertion, artificial cerebrospinal fluid (ACSF) was added to the brain’s surface, and the probes were inserted at a speed of 50µm/min to a depth of 950µm. Fifteen minutes after insertion, data acquisition was initiated as previously described ^8,98^.

All data analysis was performed using Python code written in our lab. Raw data was acquired at 20 kHz, passed through a low band pass filter at 300 Hz, and then down-sampled to 1 kHz. The noise was removed from raw traces using a 60 Hz notch filter. LFP traces were selected for each mouse from the channel (from the three shanks) showing the strongest response post visual stimulus (0.5-1s) and averaged across 20 trials for comparison between groups. These trial-averaged LFP traces were used to perform time-frequency analysis filtered at various frequencies between 4-80 Hz using the Morlet wavelet method. Power spectrum analyses were done by averaging the extracted powers normalized to baseline period dB within a time window of 0.7-1.25s post visual stimulus, and further quantified by calculating the area under the curve for these time window averaged power values for 4-8 Hz, 8-12 Hz, 12-30 Hz, 30-40 Hz, 50-70 Hz, and 30-70 Hz ranges of frequencies. Single units were obtained from raw data bandpass filtered at 300-6000 Hz using Kilosort for spike clustering. Noisy units were removed by manually sorting through Phy. Peristimulus time histograms (PSTHs) were calculated with 10 ms bins for each unit. The histogram was smoothed using a 30 ms standard deviation Gaussian kernel. The firing rates of each unit were normalized by calculating the Z-scores ((FRt-mean(FRt0-t1))/std(FRt0-t1)). Population Z-score line plots were normalized to the baseline period (0-0.5 s). Duration of oscillations was quantified using a peak detection function with a minimum peak height of at least 1.0 SD from baseline and a minimum separation of 100 ms between peaks.

Statistical analysis was done using Python packages SciPy, and Pingouin. Unit population distributions for the duration of oscillations were compared between groups by performing two-sample Kolmogorov-Smirnov tests. Shapiro-Wilk test was used to check for the normality of data. For normally distributed data, two-way ANOVA was used, followed by the corresponding Tukey-HSD test. For non-normal distributions, the Mann-Whitney U test was used.

### Directed information analysis

We conducted a directed information analysis as described previously ^9,37,38^.

#### Spike Train Preprocessing

For each unit, we used data that was between 500 ms and 1500 ms of each stimulus onset. This window was chosen to capture activity within the stimulus-induced persistent oscillatory period. Time was discretized to 1 ms bins, leading to binary-valued spike trains. Periods of time for which no units had activity for 100 ms were removed to avoid a large data imbalance.

#### Regression – Overview

With each unit’s spiking represented as a binary-valued vector, we modeled the propensity of spiking through a logistic regression model (equivalently, a generalized linear model using the logit link function). The spiking propensity could depend on each unit’s recent spiking activity as well as that of other units. For each unit *Y*, the likelihood of a spike at time t, denoted as *Y*(*t*) = 1, conditioned on a column vector *x* of exogenous variables we will describe in more detail shortly (including a constant offset) and with a row vector *θ* of model parameters, was modeled as

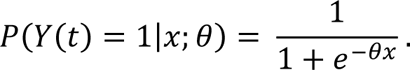

#### Regression – Exogenous Variables

In modeling the activity of each unit *Y*, *Y*’s own recent spiking activity was represented with two exogenous variables. The first variable was *Y*(*t* − 1) + *Y*(*t* − 2), designed to represent the refractory period. The second variable was *Y*(*t* − 3) + ⋯ + *Y*(*t* − 10) designed to capture self-dependence. For each other unit *X*, a single variable *X*(*t* − 1) + ⋯ + *X*(*t* − 10) was used to model *Y*’s activity. Like in ^9,33^, a 10 ms window was chosen to balance capturing effects from putative pre-synaptic units with the effects of latent common inputs.

#### Regression – Training/Testing sets

For each unit *Y*, after exogenous variables were computed, the data from trials with the same stimulus were pooled and then randomly split 50/50 into a training set and a test set. The random split was stratified by the binary *Y*(*t*) values.

#### Regression – Regularization

We used an *L*^1^ regularization term in our logistic regression (similar to the LASSO method for linear regression) to mitigate overfitting. For a parameter vector *θ*, its *L*^1^ norm, denoted as |*θ*|_1_, is the sum of the absolute values of each element except for the constant offset. Thus, the time-averaged negative log-likelihood of unit *Y*’s activity, using model vector *θ*, denoting the vector of exogenous variables computed for *Y* at time *t* as *x*(*t*), and with a hyper-parameter *λ*, was

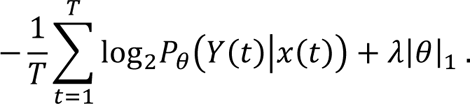

Conceptually, minimizing the first term alone, the negative log-likelihood, will lead to a model (parameterized by *θ*) that fits the data as well as possible, likely using all exogenous variables (equivalently, all elements of *θ* will be non-zero). Minimizing the second term alone, which can be viewed as a measure of model complexity, will drive elements of *θ* to 0, leading to a model that ignores all exogenous variables except the constant offset. We used a cross-validation procedure, described below, to choose the hyper-parameter *λ*.

#### Regression – Cross-validation

We used 5-fold cross-validation to select a hyper-parameter λ used for each unit *Y*. We considered forty candidate values of *λ* uniformly spaced on a log-scale between 1e-4 and 1e+4. Those values of *λ* were chosen to ensure the empirically selected *λ* was always within the largest and smallest candidate values and to balance computational complexity (considering more candidate *λ* values require more computation) with accuracy (so the selected *λ* would be close to what would have been chosen with a finer grid).

For that procedure, the training set was randomly partitioned into five parts. For each value of *λ*, each part was held out once, a model was fit on data from the remaining four parts using the current *λ*, and the log-likelihood of the resulting model was evaluated on the held-out data. The five log-likelihoods (one for each part that was held out) for the current *λ* were averaged. The hyper-parameter *λ* with the highest log-likelihood was selected. A new model was fit on all training data using that *λ*. Putative pre-synaptic units of *Y* were those with non-zero entries in the final fitted *θ* vector. We will later discuss calculating directed information from test data using the final model.

#### Regression – Implementation

Regularized regression was performed in Python (v3.8.8) using the LogisticRegressionCV function in the scikit-learn (v1.1.1) package with an ’l1’ penalty, five-folds, and with fixed hyper-parameter *λ* values for cross-validation noted above.

#### Directed Information Calculation: Overview

We measured the ‘strength’ of putative pre-synaptic connections (for each post-synaptic unit *Y*, those units *X* whose corresponding parameter in the parameter vector *θ* was non-zero) using directed information ^99^. It can be viewed as a non-parametric generalization of Granger causality ^100^. For time-series *Y*(*t*) and (multi-variate) *X*(*t*) with a joint distribution *P*(*X*(1), …, *X*(*T*), *Y*(1), …, *Y*(*T*)), if *Y* is Markov order one, the time-averaged directed information from *X* to *Y* is

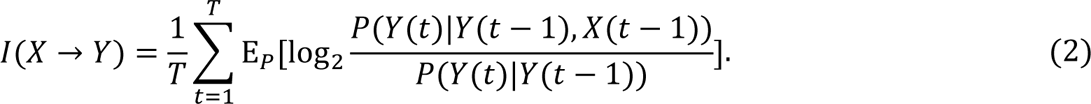

We estimated (Equation 2) by fitting models as described above and computing the time-averaged log-likelihood ratios over the test data. We then normalized directed information values by the marginal entropy

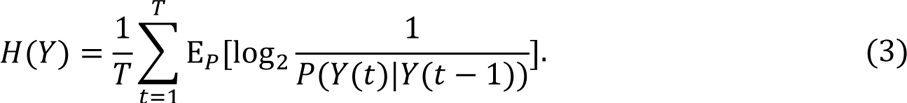

The normalized directed information *I*(*X* → *Y*)/ *H*(*Y*) can be viewed as the fraction of Y’s uncertainty that is “explained” using *X*’s past.

#### Directed Information – Area-wise Assessment

We next describe how we aggregated unit-level normalized directed information values into area-wise values. For each unit *Y*, we fit models and then evaluated directed information with candidate pre-synaptic units for each combination of recording areas: using candidate units from both V1 and LM, only using units from V1, only using units from LM, and a model only using *Y*’s own past. Let *θ*_All_, *θ*_*V*1_, *θ*_AL_, and *θ_None_* denote the corresponding model parameter vectors and let *I*_All_, *I*_*V*1_, *I*_*V*1_, and *I_None_* = 0 denote the respective directed information values.

We attribute the marginal contributions of putative pre-synaptic units from each area to a unit *Y* using Shapley values ^101^. Shapley values are a game-theoretic solution for fairly dividing a reward in collaborative games based on each members’ average (over all subsets of other members) marginal contribution. For two areas, the formula is

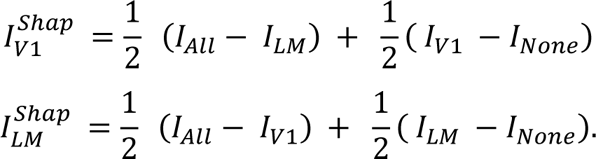

For V1 and LM, we calculated the median of the Shapley values over each unit *Y* for each experimental setting (pre/post-training, WT/FX). We then calculated the differences between these median values for post and pre-training.

#### Significance Testing

We computed the significance of differences post and pre-training using Monte Carlo approximations for two-sided permutation tests. 1e+5 Monte Carlo iterations were used for each difference reported. For each iteration, each Shapley value (of normalized directed information values) used to calculate the median values from a pre-synaptic area to a unit in a post-synaptic area was relabeled as pre-training or a post-training connection using corresponding proportions of connections in the pre- and post-stimulus training experiments. The difference of the medians (after relabeling) was then computed.

The Monte Carlo approximations were computed in Python (v3.8.8). We corrected the p-values separately for each Figure with the Benjamini-Hochberg procedure (Benjamini and Hochberg, 1995), implemented in the Python statsmodels (v0.13.2) package.

### Acute V1 slices preparation

The naïve or visually experienced mice with virus injections were anesthetized by the mix of saline-diluted ketamine (0.1mg/g body weight) and xylazine (0.16mg/g body weight), and deep anesthesia was confirmed by toe pinching. The perfusion and acute slice recovery strategies were the same as described in the previous paper of our lab ^9^ except for substituting sucrose holding cutting buffer to choline holding buffer. The mice were first perfused with ice-cold choline artificial cerebrospinal fluid (choline ACSF, composition in mM: 110 NMDG, 30 NaHCO3, 25 Dextrose, 11.6 ascorbic acid, 7 MgCl2, 3.1 Na-pyruvate, 2.5 KCl, 0.5 CaCl2; pH=7.3-7.4) transcardially till livers turned pale. The mice were then decapitated, and the brains were extracted and trimmed to be glued to the cutting chamber of the vibratome (VT 1000 Leica). 300-micrometer coronal brain sections containing the V1 were cut in ice-cold carbogenating (carbogen: 95% O2, and 5% CO2). choline ACSF by vibratome. The slices were then transferred to an incubation chamber warmed up in 32 °C water bath filled with ACSF (composition in mM: 124 NaCl, 26 NaHCO3, 10 Dextrose, 2.5 KCl, 2 CaCl2, 1.25 NaH2PO4, 0.8 MgCl2; pH=7.3-7.4) for 30 min. The chamber was removed from the water bath and slices were incubated at room temperature for one hour. After the equilibrium, the slices were ready for clamp-patch recording, and the slices were healthy for 10 hours in the incubation chambers.

### Electrophysiology and data acquisition for whole-cell patch-clamp recording

Patch-clamp recordings were conducted on an electrophysiology rig (SliceScope Pro 1000, Scientifica). The brain slices were settled down in a submersion-type recording chamber with a slice hold down and perfused with carbogenated ACSF (composition in mM: 124 NaCl, 26 NaHCO3, 10 Dextrose, 2.5 KCl, 2 CaCl2, 1.25 NaH2PO4, 0.8 MgCl2; pH=7.3-7.4) continuously during the whole recordings. Neurons were imaged with infrared differential interference contrast optics from the rig equipped with a 4x objective lens and a 40x water immersion objective lens (Olympus) and ROLERA™ Bolt Scientific CMOS Camera (QImaging). The resistance of the recording pipette was 4-7 MO, and the intracellular pipette solution (composition in mM: 103 D-gluconic acid, 103 CsOH, 20 HEPES, 10 Na2-phosphocreatine, 5 QX314-Cl, 5 TEA-Cl, 4 Mg-ATP, 2.8 NaCl, 0.3 Na2-GTP, 0.2 EGTA; pH=7.2-7.3; 290 mOsm). In some experiments, a small amount of 4% w/v Alexa Fluor 594 (A-10438, ThermoFisher Scientific) dissolved in internal solution was back-loaded into the glass electrode to label the patched cell. All neurons were whole-cell patched in voltage-clamp mode with a holding potential of -70 mV. All EPSCs were recorded in voltage-clamp mode with a holding potential of -80 to -70 mV while IPSCs were recorded at 0 to 10 mV. Electrophysiological data were acquired with an amplifier (Multiclamp 700B, Molecular Devices) and a digitizer (Digitata, 1550, Molecular Devices). The acquired data were recorded with Clampex 10.4 (Molecular Devices) and filtered with a 20k Hz low pass filter. The evoked excitatory postsynaptic currents (EPSC) were recorded in a whole-cell voltage patch-clamp, holding at -70 mV. The membrane resistance of each patched neuron was recorded from the readout of the Clampex 10.4.

### Channelrhodopsin-Assisted Circuit Mapping (CRACM)

To measure the FF V1→LM and FB LM→V1, we patched the excitatory pyramidal neurons in LM or V1 layer 2/3, 4, 5 correspondingly, identified by the morphology and relative location in the cortical layers of coronal slices. 470 nm LED light to activate ChR2 was generated with an LED light source (High-Power LED Collimator Source, 470 nm, 50W, Mightex), and the light stimulation patterns were produced by a patterned illuminator (Polygon 400, Mightex) ^102^. The square light, 67 μm x 67 μm, was delivered in a 10 by 10 grid covering 670 μm x 670 μm areas of the cortical regions centered at patched neurons in a pseudo-random sequence through the 10x objective lens. The LED and patterned illuminator were controlled by the manufacturer’s software, and stimulation and recording were synchronized by the Digidata. CRACM heatmaps were plotted from light-induced EPSC amplitudes at each pixel.

### Histology

The acute brain slices with Alexa Fluor 568 dye-filled were fixed with 4% paraformaldehyde (PFA) and imaged by a confocal microscope (Zeiss 710). For Super-resolution imaging sample preparation, we used the male litters of male Thy1-ChR2-EYFP cross with female B6 for control and maleThy1-ChR2-EYFP cross female Fmr1 KO.

The 3 to 4-month-old mice were anesthetized by intraperitoneal injections of a mix of 90 mg/kg ketamine (59399-114-10, Akron) and 10 mg/kg xylazine (343750, HVS) and transcardially perfused with phosphate-buffered saline (1xPBS, 1:10 diluted from DSP32060, Dot Scientific), followed by 4% PFA (1:4 diluted with 1× PBS from Paraformaldehyde 16% Aqueous Solution, Electron Microscopy Sciences) to pre-fix the brain. The brain was excavated from the skull and immersed in 4% PFA at 4 °C overnight. The fixed brain sample was then cut into 30-50 μm coronal sections on a vibratome (1000 Plus, Vibratome).

For immunohistochemistry staining, the brain sections were washed in wash buffer (0.1% Triton X-100 in 1× PBS) three times with a gentle shake (120 rpm, Orbi-Shaker, Benchmark), 15 min for each time. The brain sections were then incubated in a blocking buffer (5% BSA (A9647, Sigma-Aldrich) in 1× PBS) for 1.5 h with a gentle shake. After blocking, the brain sections were incubated with chicken anti-GFP antibody (ab13970, Abcam, diluted to 1:1,000 in blocking buffer) at 4 °C overnight. To remove extra primary antibodies, the sections were washed three times as in the first step. The slices were incubated with goat anti-chicken Alexa Fluor 647-conjugated antibody (A21449, Invitrogen, diluted to 1:600 in wash buffer) for 2 h with a gentle shake. After being washed three times, the sections were kept in 1x PBS before being mounted for super-resolution imaging.

### 4Pi super-resolution microscope setup

A 4Pi single-molecule localization microscope was built using dual opposing 100x/1.4-NA oil-immersion objectives (UPLSAPO 100XO, Olympus) and an sCMOS camera (Orca-Flash4.0v2, Hamamatsu). This setup allowed for a magnification of 50x, a field of view of 20x20 μm², and an effective pixel size of 129 nm^32,60,103,104^. The coherent fluorescence from both objectives coincided and was aligned using nonpolarized and polarized beam splitters (NT47-009 and NT49-002, Edmund Optics) to generate four beams. These four beams were directed to different regions on the sCMOS camera and acquired simultaneously. Customized Babinet-Soleil BK7/Quartz wedges were used to control the P- and S-polarized phase delays, and two deformable mirrors (Multi-5.5, Boston Micromachines) were placed in conjugated pupil planes of respective objectives to control systematic aberrations. A 642-nm continuous-wave laser (2RU-VFL-P-2000-642-B1R, MPB Communications) was utilized for HILO excitation through the bottom objective. The tilt angle of the excitation was optimized to 54°, enabling the highest signal-to-background ratio during imaging. Quad-band dichroic and bandpass filters (FF01-446/523/600/677-25, Di01-R405/488/561/635-17.5×24, Semrock) were used to filter fluorescence from the excitation and unwanted background. The microscope setup was synchronized through a custom LabVIEW program.

### Optical clearing and sample mounting

Right before data acquisition, the labeled brain sections were placed on cleaned coverslips, and the visual cortex regions were positioned at the center of the coverslips. The optical clearing was performed by adding 200 μL ultrafast optical clearing (FOCM) reagent of 30% wt/vol urea (U15-500, Fisher Scientific), 20% wt/vol D-sorbitol (S1876-500G, Sigma-Aldrich), and 5% wt/vol glycerol (G5516, Sigma-Aldrich) in DMSO (276855, Sigma-Aldrich) on the samples for 1-2 minutes at room temperature^105^. After optical clearing, the samples were washed once with 1xPBS and 1 μL 200-nm crimson beads (1:10^6^ diluted from custom-designed beads in deionized water, Invitrogen) were added near the outer edge region of the visual cortex. The samples were mounted with 70 μL freshly prepared cyclooctatetraene-based buffer (10% wt/vol glucose in 50 mM Tris, 50 mM NaCl, 10 mM MEA (M9768, Sigma-Aldrich), 50 mM BME (63689, Sigma-Aldrich), 2 mM COT (138924, Sigma-Aldrich), 2.5 mM PCA (37580, Sigma-Aldrich) and 50 nM PCD (P8279-25UN, Sigma-Aldrich), pH 8.0) and another coverslip on top^106^. The sandwiched samples were sealed using silicone dental glue (Twinsil speed 22, Picodent) and left to dry for 30 minutes at room temperature.

### Spine data acquisition and analysis

The excitation power was set to 30 W/cm^2^ for finding regions of interest (ROI) and bead measurement. Before imaging, excitation was elevated to 12 kW/cm^2^ and scanned around the ROI to transfer most of the fluorophores into dark states and bleach autofluorescence for 30-60 seconds. For imaging, the excitation was set to 7 kW/cm^2^, and 100,000 to 140,000 frames of raw blinking images were recorded with an acquisition time of 20 ms. Each raw image was first cropped into four quadrants based on arranged regions on the sCMOS camera and underwent 4Pi super-resolution reconstruction using the phase-retrieval-based 4Pi single molecule localization algorithm (PR-4Pi)^104^. 4Pi point spread function (4Pi-PSF) was modeled using the two pupil functions of objectives and the interfered bead stack from a single bead near the ROI. To retrieve pupil functions, the bead was scanned across z positions from –1 to 1 µm, with a step size of 100 nm, a frame rate of 10 Hz, and 5 frames per z position, and acquired fluorescence from the top and bottom objectives, respectively. The resulting bead measurements were subjected to a phase retrieval algorithm to obtain the pupil functions. The interferometric bead data was acquired by scanning across z positions from –1 to 1 µm, with a step size of 20 nm, a frame rate of 10 Hz, and 5 frames per z position, using interfered coherent fluorescence.

### 3D Spine structure quantification

The targeted spine was manually identified from the reconstruction results by marking along the spine neck to the spine head. The software then traced these marks as a curve (’fit’, model type ’smoothingspline’, MATLAB), straightened the spine along the X direction, and segmented the localizations into cross sections. Each cross-section possessed dimensions of 600-1000 nm in width, 600-1000 nm in height, and 150 nm in length, with a 50 nm overlap in X direction between neighbors. For visualization, the segmented localizations were rendered as XY and YZ cross-sectional images with a pixel size of 6 nm. These images were pseudo-colored based on their relative Z positions and were smoothed using a 2D Gaussian blur with an RMS width of 6 nm. To quantify the structure, the software captured the boundaries of the largest structures from discrete localizations (’alphaShape’, alpha shape radius of 0.4, MATLAB). The structural centers and the semi-minor axis of the structures were estimated by fitting an elliptical function to the localizations within the alpha-shape area, which were used in the spine area and volume calculation.

