## Supplemental Figure S1-11 for "Impaired Experience-Dependent Theta Oscillation Synchronization and Inter-Areal Synaptic Connectivity in the Visual Cortex of Fmr1 KO Mice"

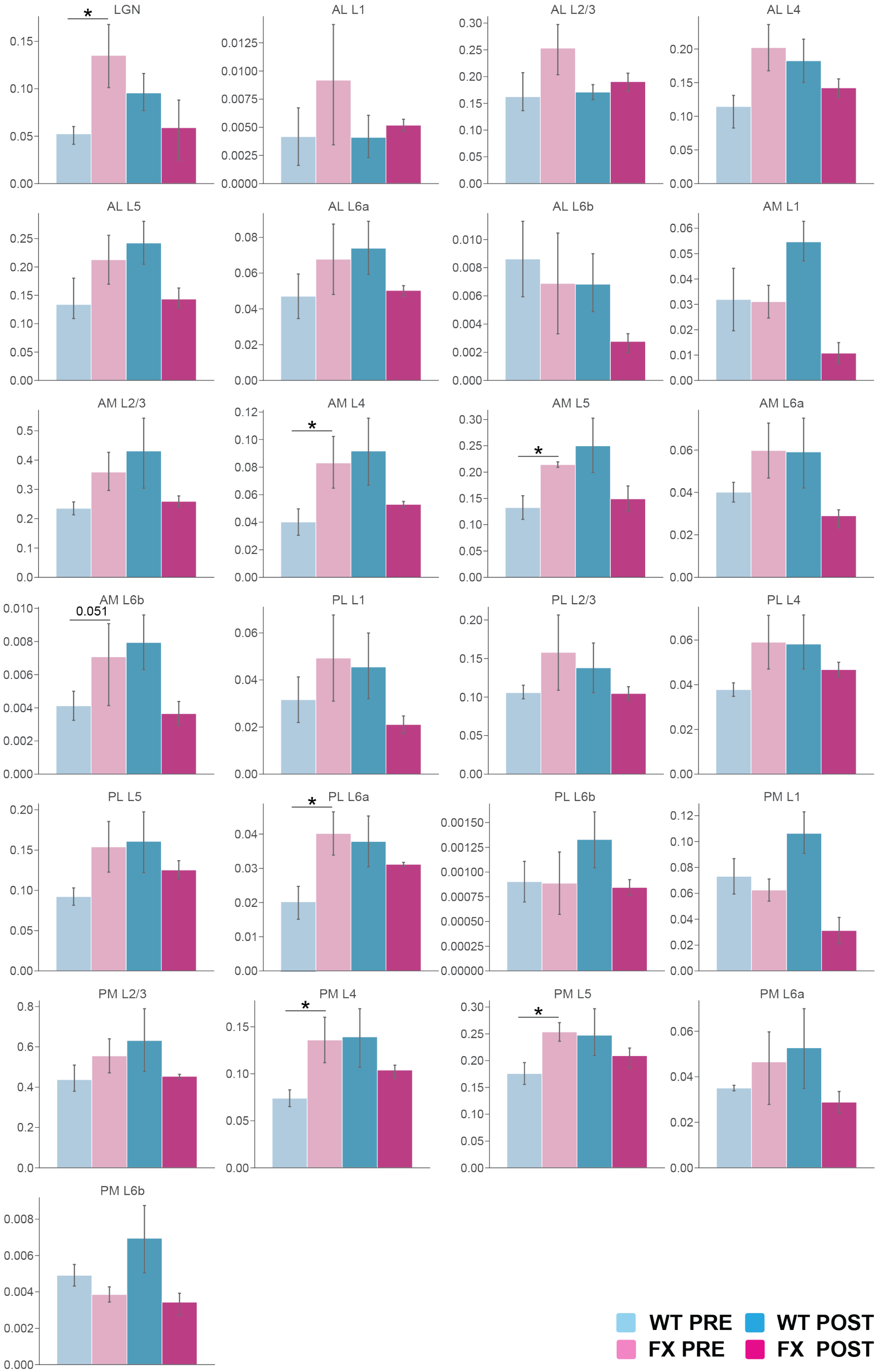
LGN
AL L1
AL L2/3
AL L4
AL L5
AL L6a
AL L6b
AM L1
AM L2/3
AM L4
AM L5
AM L6a
AM L6b
0.051
PL L1
PL L2/3
PL L4
PL L5
PL L6a
PL L6b
PM L1
PM L2/3
PM L4
PM L5
PM L6a
PM L6b
WT PRE
WT POST
FX PRE
FX POST

**Figure S1. Normalized C-fos+ expression in visual areas other than V1 and LM, related to Figure 1.**

The significance was reported by the T-test (p-values in Table X).

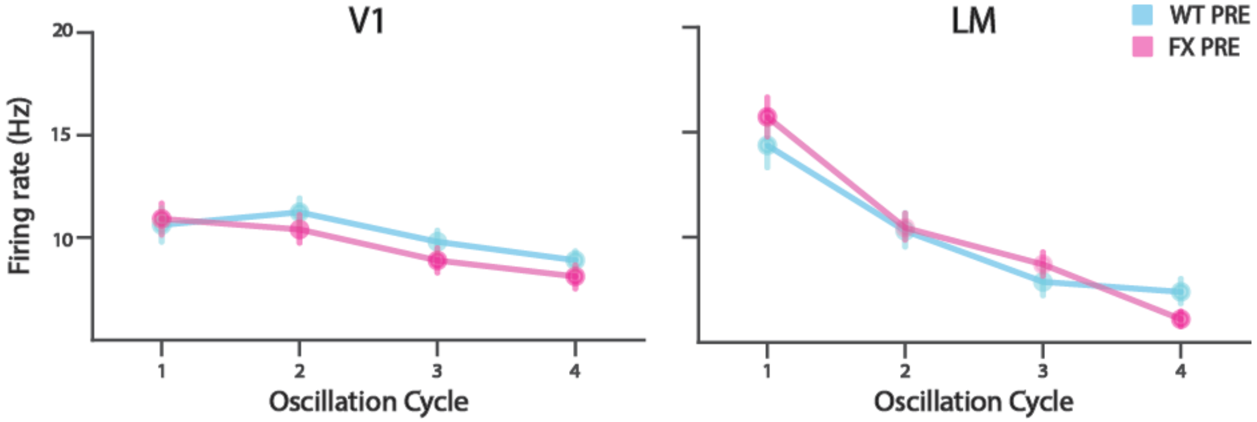


**Figure S2. Unit population firing rates pre-training for V1 and LM, related to Figure 3**

Left: Maximum firing rates at 4 oscillation cycle time windows averaged across units in V1 pre-training (WT: N=267 units, n=10 mice, FX: N=270 units, n=8 mice) in WT (cyan) and FX (magenta) mice. Right: Maximum firing rates at 4 oscillation cycle time windows averaged across units in LM pre training (WT: N=322 units, n=10 mice, FX: N=410 units, n=8 mice) in WT (cyan) and FX (magenta) mice.

*$p<0.05$.

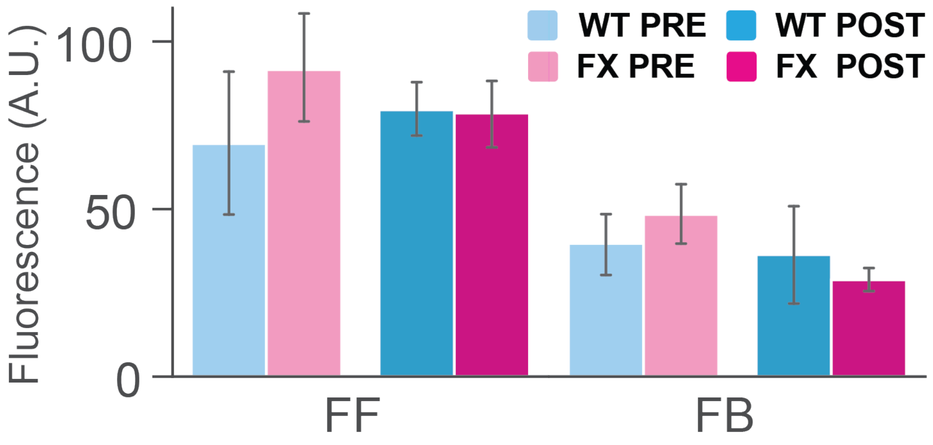


**Figure S3. Channelrhodopsin 2 expression level in the patched region for CRACM measured by fluorescence of EYFP, related to Figure 4 and 5.**

Bar graphs of average fluorescence of EYFP in the patched region in feedforward projection measurement (WT pre: N=7, WT post: N=8, FX pre: N=6, FX post: N=12 fields of view). Significance was reported from two-way ANOVA followed by Tukey's HSD tests (two-way ANOVA: genotype: $F = 0.33$; $p = 0.57$. Perceptual experience: $F = 0.0155$; $p = 0.90$. Tukey's post hoc: WT pre versus FX pre: p = 0.46; WT post versus FX post: p = 0.9; WT pre versus WT post: p = 0.67; FX pre versus FX post: p = 0.51).

Bar graphs of average fluorescence of EYFP in the patched region in feedback projection measurement (WT pre: N=12, WT post: N=6, FX pre: N=9, FX post: N=8 fields of view). Significance was reported from two-way ANOVA followed by Tukey's HSD tests (two-way ANOVA: genotype: F = 3.75E-3; p = 0.95. Perceptual experience: F = 1.43; p = 0.24. Tukey's post hoc: WT pre versus FX pre: p = 0.52; WT post versus FX post: p = 0.59; WT pre versus WT post: p = 0.85; FX pre versus FX post: p = 0.082).

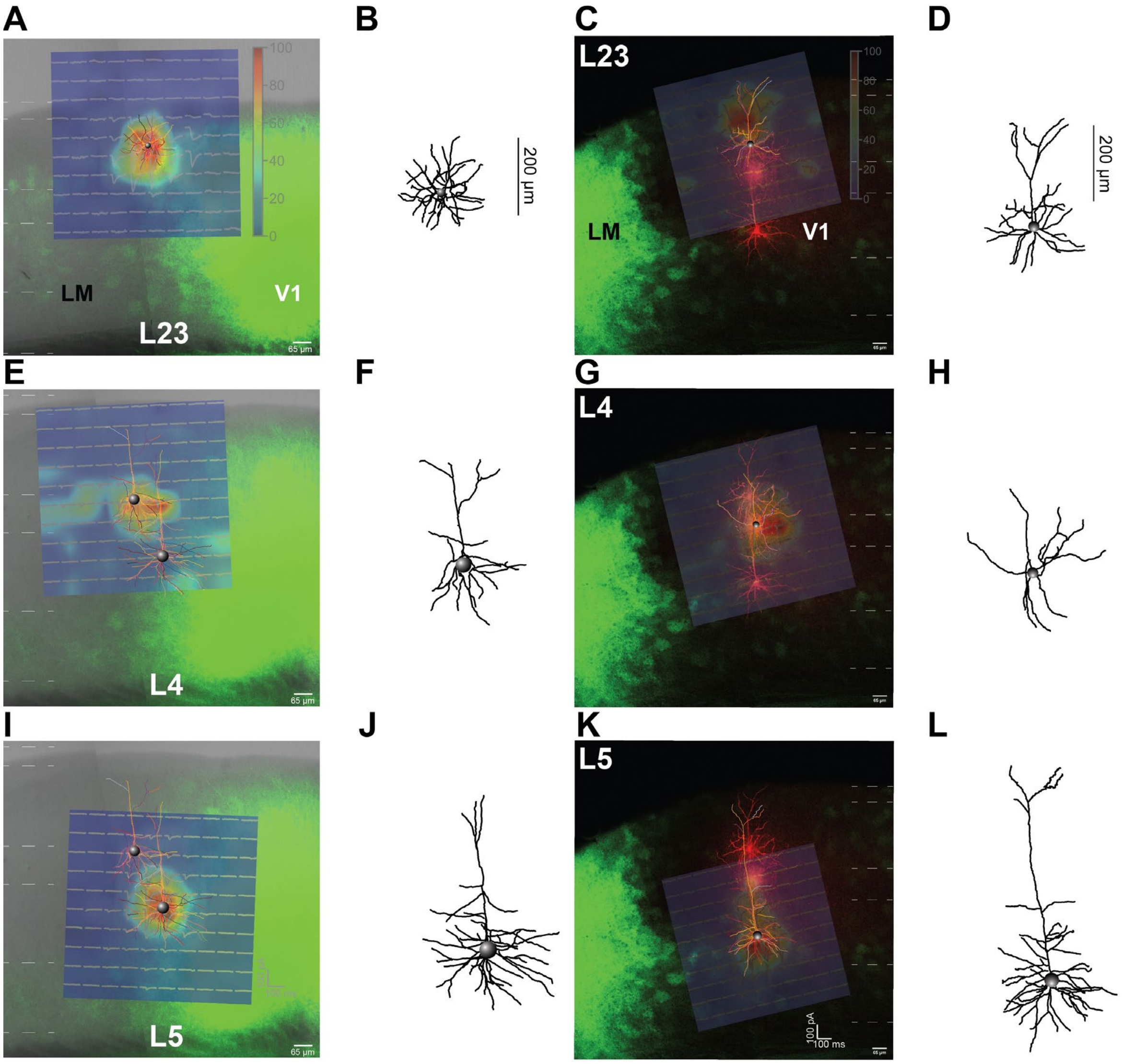


**Figure S4. Examples of cell trace and corresponding CRACM maps from each layer of LM, related Figure 4 and 5.**

(A) Example of a tiled composite confocal image of mapped LM L23 neuron (red) filled with Alexa Fluor 568 Hydrazide (scale bar, 65μm). The green fluorescence indicates ChR2-YFP-positive neurons. The corresponding sCRACM map (10×10 grids, 65um space) with $CRACM_{EPSC}$ traces was overlaid with the filled neurons.
(B) Traced PC from (A) showing the morphology of a patched neuron of L23.
(C) Same as (A) but for V1 L2/3 neuron.
(D) Same as (B) but for V1 L2/3 neuron.
(E) Same as (A) but for LM L4 neuron.
(F) Same as (B) but for LM L5 neuron.

(G) Same as (A) but for V1 L4 neuron.

(H) Same as (B) but for V1 L4 neuron.
(I) Same as (A) but for LM L5 neuron.
(J) Same as (B) but for LM L5 PC.
(K) Same as (A) but for V1 L5 PC.
(L) Same as (B) but for V1 L5 PC.

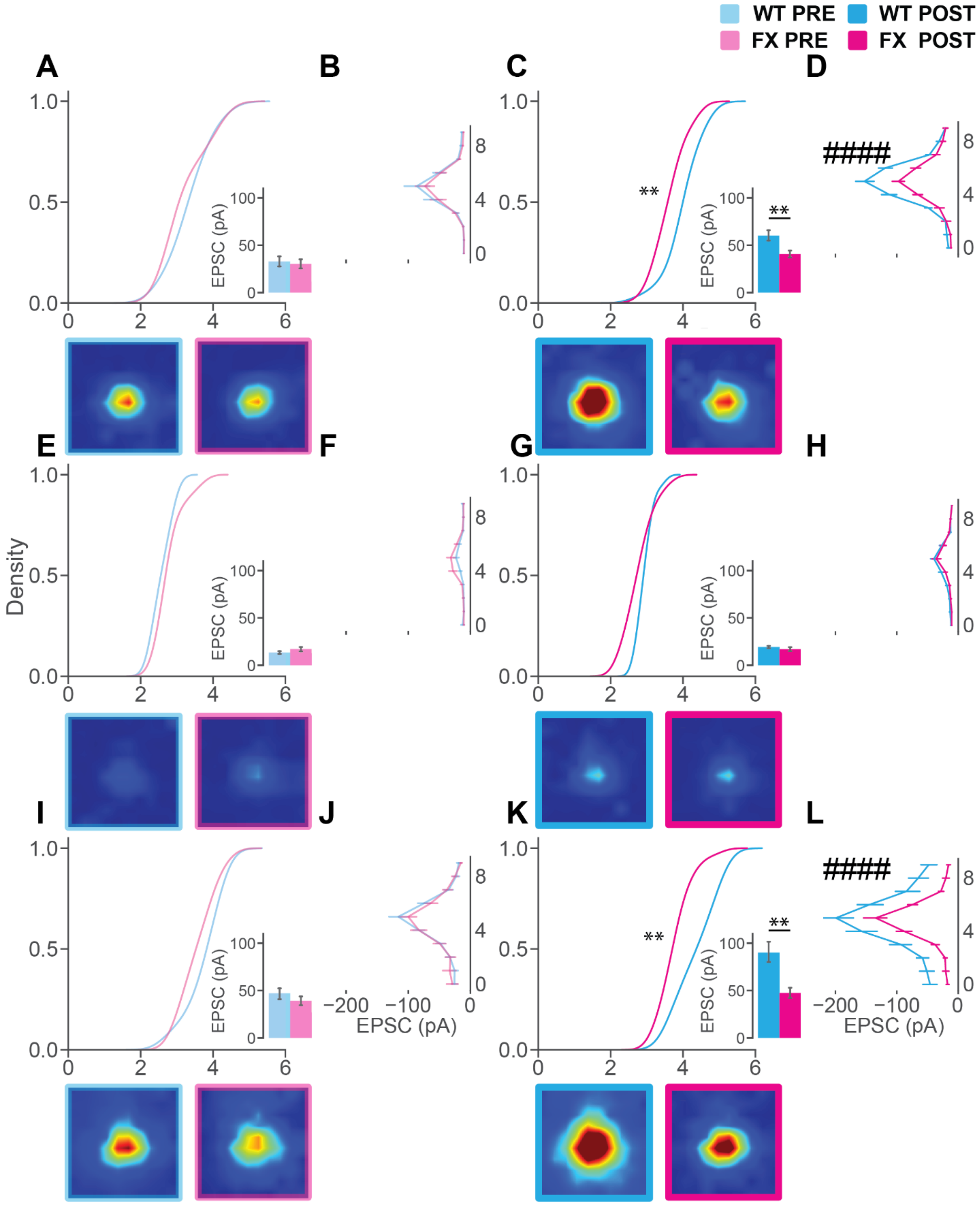


**Figure S5. FF sCRACM maps of input from V1 ChR2 positive neurons to PCs in LM comparing between WT and FX groups, related Figure 4.**

**(A)** The cumulative distributions (top) of the averaged $EPSC_{CRACM}$ amplitudes plotted in ln scale in sCRACM map recorded from LM L23 PCs. Significance was reported from Kolmogorov–Smirnov test (p= 0.315, WT naïve L23: 15 cells, 6 mice L23; FX naïve: 23 cells, 6 mice). Bar graphs of averaged $EPSC_{CRACM}$ amplitudes ± SEM for WT pre- and post-training groups (inset). Significance was reported from the Mann–Whitney U test (p= 0.420). The average sCRACM maps across each grid were plotted below the corresponding cumulative.

**(B)** Averaged $EPSC_{CRACM}$ amplitudes ± SEM by grid position in the vertical direction (perpendicular to the brain surface, top) from V1 L2/3 PCs in naïve mice. Significance was reported from two-way ANOVA (genotype: F = 0.626; p = 0.429). Averaged $EPSC_{CRACM}$ amplitudes ± SEM by grid position in the tangential direction (parallel to the brain surface, bottom). Significance was reported from two-way ANOVA (genotype: F = 0.760; p = 0.384).

**(C)** Same as (A), but from post-training groups. Significance was reported from Kolmogorov–Smirnov test (p= 4.195E-3, WT post-training L23: 28 cells, 8 mice; FX post L23: 30 cells, 8 mice). Significance was reported from the Mann–Whitney U test (p= 5.181E-3).

**(D)** Same as (B), but from post-training groups. In the vertical direction, significance was reported from two-way ANOVA (genotype: F = 36.47; p = 2.824E-9). In the tangential direction, significance was reported from two-way ANOVA (genotype: F = 41.51; p = 2.524E-10).

**(E)** Same as (A), but from L4 PCs. Significance was reported from Kolmogorov–Smirnov test (p= 0.468, WT naïve L4: 6 cells, 5 mice; FX naïve L4: 12 cells, 6 mice). Significance was reported from Mann–Whitney U test (p= 0.494).

**(F)** Same as (B), but from L4 PCs. In the vertical direction, significance was reported from two-way ANOVA (genotype: F = 2.078; p = 0.151).

**(G)** Same as (C), but from L4 PCs. Significance was reported from Kolmogorov–Smirnov test (p= 0.146, WT post-training L4: 15 cells, 8 mice; FX post-training L4: 13 cells, 8 mice). Significance was reported from the Mann–Whitney U test (p= 0.128).

**(H)** Same as (D), but from L4 PCs. In the vertical direction, significance was reported from two-way ANOVA (genotype: F = 2.450; p = 0.119).

**(I)** Same as (A), but from L5 PCs. Significance was reported from Kolmogorov–Smirnov test (p= 0.191, WT naïve L5: 12 cells, 6 mice; FX naïve L5: 18 cells, 6 mice). Significance was reported from Mann–Whitney U test (p= 0.197).

**(J)** Same as (B), but from L5 PCs. In the vertical direction, significance was reported from two-way ANOVA (genotype: F = 0.0987; p = 0.754).

**(K)** Same as (C), but from L5 PCs. Significance was reported from Kolmogorov–Smirnov test (p= 2.408E-3, WT post-training L5: 21 cells, 8 mice; FX post-training L5: 26 cells, 8 mice). Significance was reported from the Mann–Whitney U test (p= 1.720E-3).

**(L)** Same as (D), but from L5 PCs. In the vertical direction, significance was reported from two-way ANOVA (genotype: F = 63.16; p = 1.549E-14). Data were presented as mean ± SEM. Two-way ANOVA: #p<0.05, ##p<0.01, ###p<0.001, ####p<0.0001. Kolmogorov–Smirnov test and Mann–Whitney U test: *p<0.05, **p<0.01, ***p<0.001, ****p<0.0001

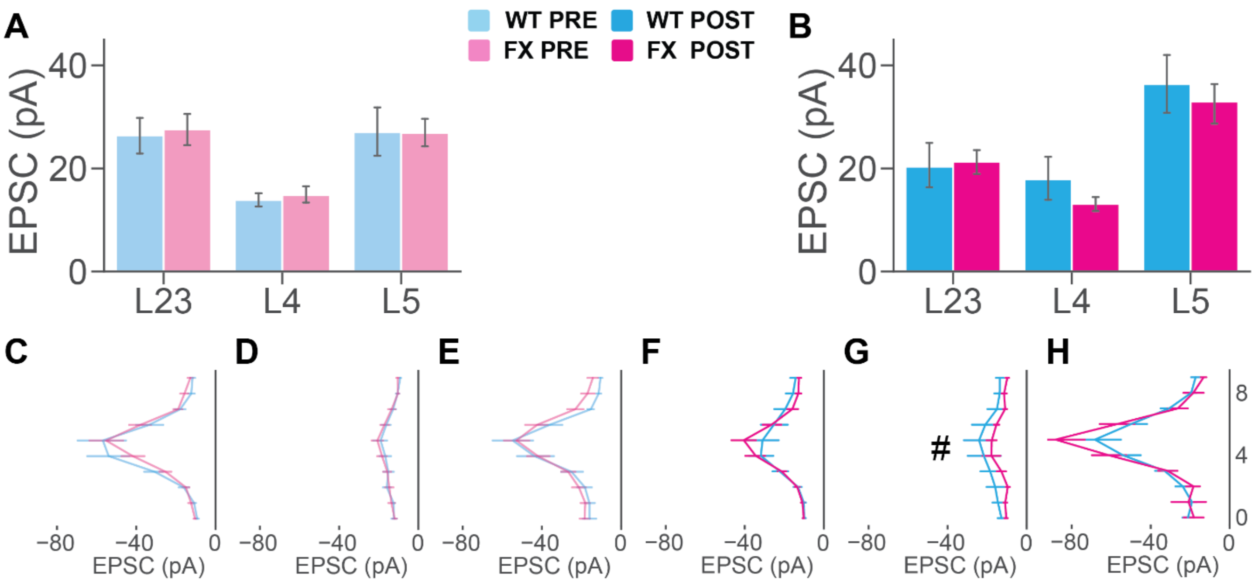


**Figure S6. FB sCRACM maps of input from LM ChR2 positive neurons to PCs in V1 comparing between WT and FX groups, related Figure 5.**

**(A)** Bar graphs of averaged EPSCCRACM amplitudes ± SEM for WT and FX pre-training groups. Significance was reported from the Mann–Whitney U test (WT naïve L2/3: 21 cells, 10 mice; FX pre L2/3: 14 cells, 5 mice; p= 0.945. WT naïve L4: 21 cells, 7 mice; FX pre L4: 11 cells, 5 mice; p= 0.925. WT naïve L5: 21 cells, 8 mice L5; FX pre: 35 cells, 13 mice; p= 0.310).

**(B)** Bar graphs of averaged $EPSC_{CRACM}$ amplitudes ± SEM for WT and FX post groups. Significance was reported from the Mann–Whitney U test (WT post-training L2/3: 17 cells, 7 mice; FX post-training L2/3: 26 cells, 11 mice; p= 0.378. WT post L4: 7 cells, 7 mice; FX post L4: 7 cells, 7 mice; p= 0.902. WT post-training L5: 17 cells, 8 mice L5; FX post: 20 cells, 13 mice; p= 0.819).

**(C)** Averaged $EPSC_{CRACM}$ amplitudes ± SEM by grid position in the vertical direction (perpendicular to the brain surface, top). Significance was reported from two-way ANOVA (genotype: F = 0.122; p = 0.727) from V1 L2/3 PCs in naïve mice

**(D)** Same as (R), but from L4 PCs. In the vertical direction, significance was reported from two-way ANOVA (genotype: F = 0.382; p = 0.537).

**(E)** Same as (R), but from L5 PCs. In the vertical direction, significance was reported from two-way ANOVA (genotype: F = 0.122; p = 0.727).

**(F)** Same as (R), but from post-training groups. In the vertical direction, significance was reported from two-way ANOVA (genotype: F = 0.0935; p = 0.760).

**(G)** Same as (U), but from L4 PCs. In the vertical direction, significance was reported from two-way ANOVA (genotype: F = 5.862; p = 0.0170).

**(H)** Same as (U), but from L5 PCs. In the vertical direction, significance was reported from two-way ANOVA (genotype: F = 0.250; p = 0.617).

Data were presented as mean ± SEM. Two-way ANOVA: #p<0.05, ##p<0.01, ###p<0.001, ####p<0.0001. Kolmogorov–Smirnov test and Mann–Whitney U test: *p<0.05, **p<0.01, ***p<0.001, ****p<0.0001

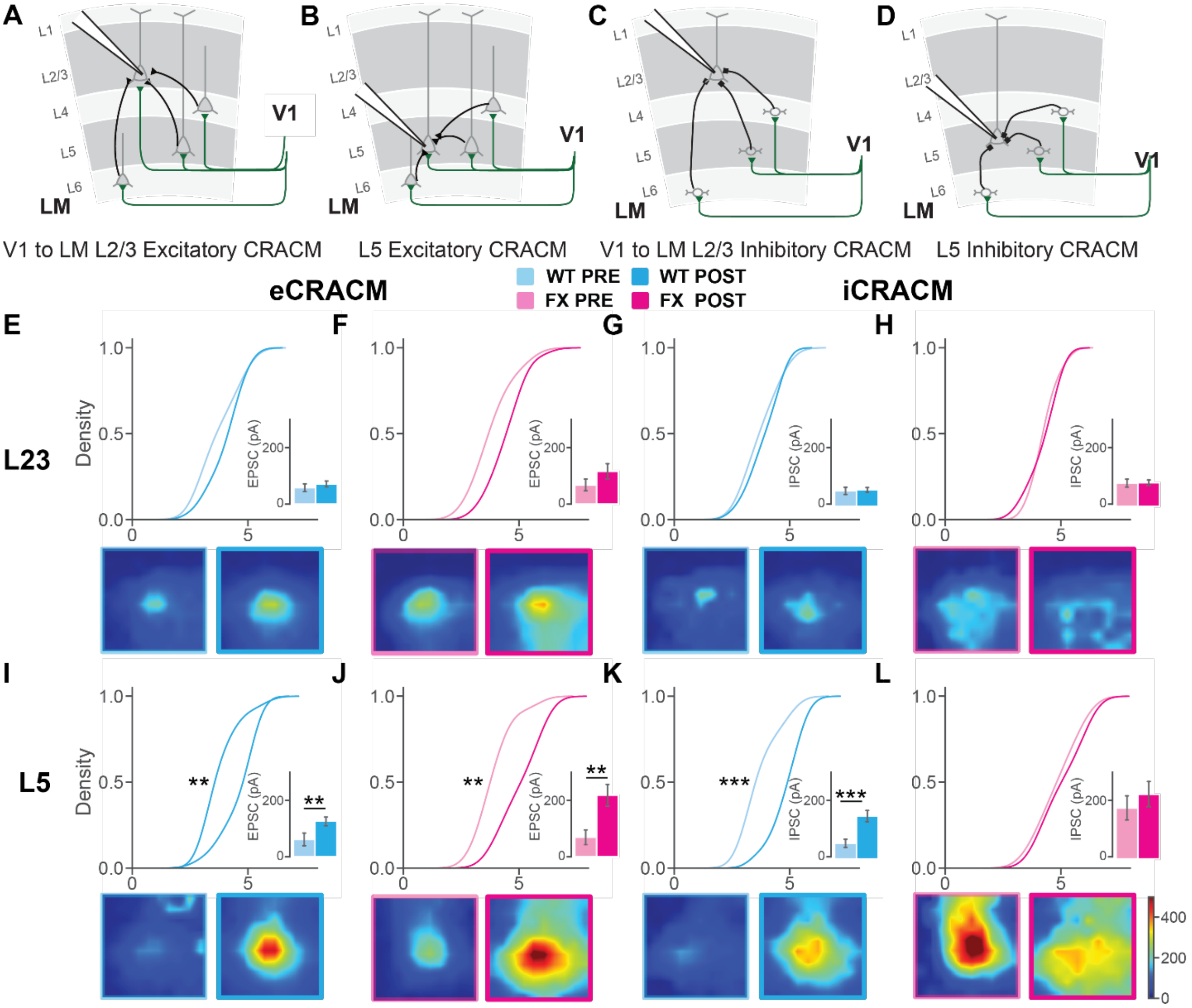


**Figure S7. Multi-synaptic excitatory and inhibitory FF CRACM maps of input from V1 ChR2 positive neurons to PCs in LM, related Figure 4.**

**(A)** Illustration of V1 to LM L2/3 PCs feedforward excitatory projections.
**(B)** Same as (A) but to L5 PCs.
**(C)** Illustration of V1 to LM L2/3 PCs feedforward inhibitory projections.
**(D)** Same as (B) but to L5 PCs.
**(E)** The cumulative distributions (top) of the averaged $EPSC_{CRACM}$ amplitudes plotted in ln scale in CRACM map recorded from LM L2/3 PCs. Significance was reported from Kolmogorov–Smirnov test (p= 0.427, WT naïve L23: 11 cells, L23; WT post: 18 cells). Bar graphs of averaged $EPSC_{CRACM}$ amplitudes ± SEM for WT pre and post-training groups (inset). Significance was reported from the Mann–Whitney U test (p= 0.334). The average CRACM maps across each grid were plotted below the corresponding cumulative.
**(F)** Same as (E) but from FX pre and post-training mice groups (Kolmogorov–Smirnov test: p= 0.115, Mann–Whitney U test: p= 0.0655; WT naïve L23: 11 cells, L23; WT post: 18 cells).

**(G)** Same as (E) but for IPSCs. (Kolmogorov–Smirnov test: p= 0.752, Mann–Whitney U test: p= 0.486; WT naïve L23: 11 cells, L23; WT post: 18 cells).

**(H)** Same as (F) but for IPSCs. (Kolmogorov–Smirnov test: p= 0.658, Mann–Whitney U test: p= 0.788; WT naïve L23: 11 cells, L23; WT post: 18 cells).

**(I)** same as (E) but from FX pre- and post-training mice groups. (Kolmogorov–Smirnov test: p= 3.81E-3, Mann–Whitney U test: p= 4.5E-3; WT naïve L23: 11 cells, L23; WT post: 18 cells).

**(J)** Same as (I) but from FX pre- and post-training mice groups. (Kolmogorov–Smirnov test: p= 9.71E-3, Mann–Whitney U test: p= 3.25E-3; WT naïve L23: 11 cells, L23; WT post: 18 cells).

**(K)** Same as (I) but from FX pre- and post-training mice groups. (Kolmogorov–Smirnov test: p= 8.66E-4, Mann–Whitney U test: p= 8.43E-4; WT naïve L23: 11 cells, L23; WT post: 18 cells).

**(L)** Same as (I) but from FX pre- and post-training mice groups. (Kolmogorov–Smirnov test: p= 0.918, Mann–Whitney U test: p= 0.485; WT naïve L23: 11 cells, L23; WT post: 18 cells).

Error bars indicate SEM. $*p < 0.05$; $**p < 0.01$; $***p < 0.001$.

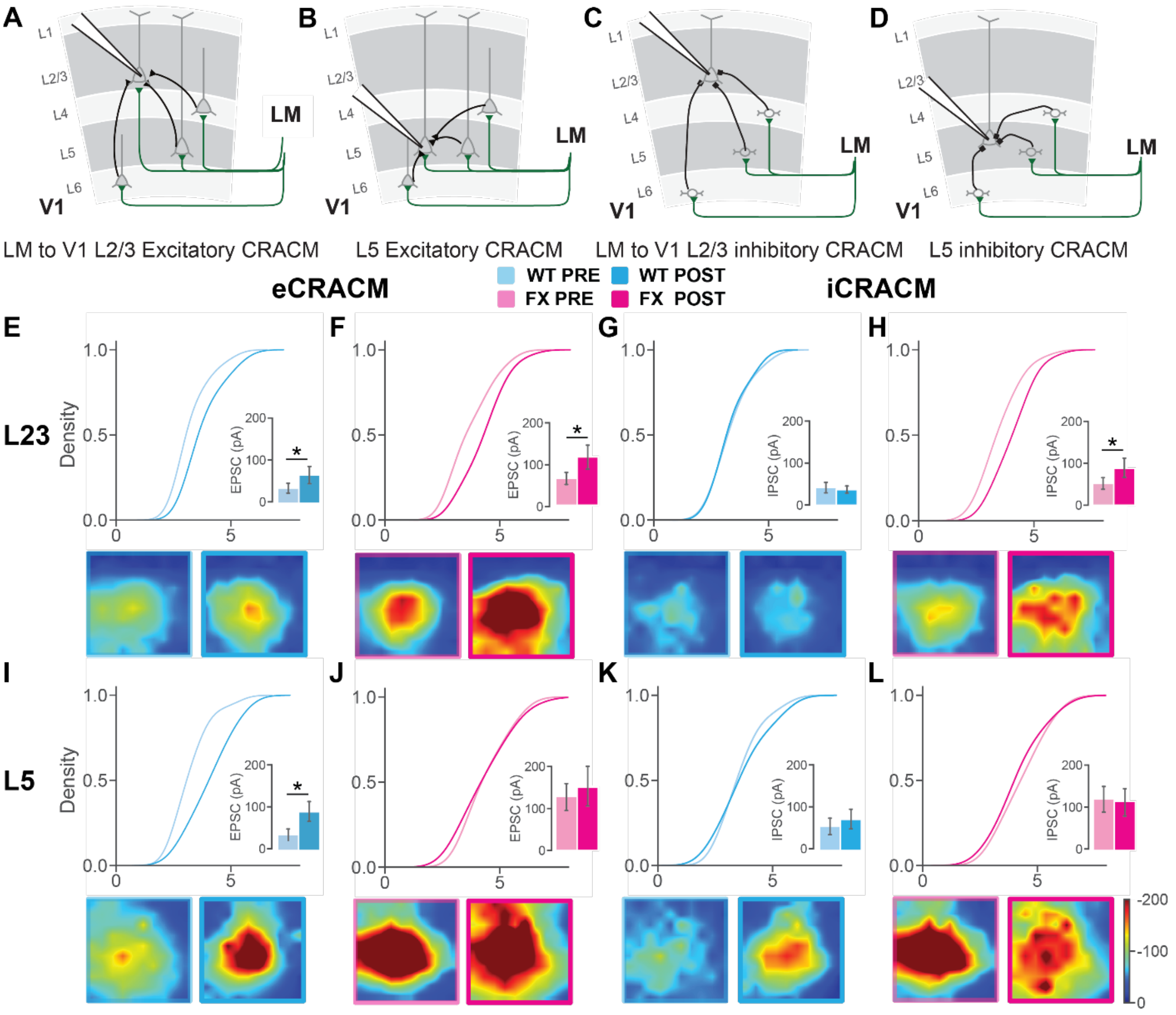


**Figure S8. Multi-synaptic excitatory and inhibitory FB CRACM maps of input from LM ChR2 positive neurons to PCs in V1, related to Figure 5.**

**(A)** Illustration of V1 to LM L2/3 PCs feedforward excitatory projections.

**(B)** Same as (A) but to L5 PCs.

**(C)** Illustration of V1 to LM L2/3 PCs feedforward inhibitory projections.

**(D)** Same as (B) but to L5 PCs.

**(E)** The cumulative distributions (top) of the averaged $EPSC_{CRACM}$ amplitudes plotted in ln scale in sCRACM map recorded from LM L2/3 PCs. Significance was reported from Kolmogorov–Smirnov test (p= 0.081, WT naïve: 14 cells, 6 mice; WT post: 15 cells, 7 mice). Bar graphs of averaged $EPSC_{CRACM}$ amplitudes ± SEM for WT pre and post-training groups (inset). Significance was reported from the Mann–Whitney U test (p= 0.042). The average eCRACM maps across each grid were plotted below the corresponding cumulative.

**(F)** Same as (E) but from FX pre and post-training mice groups (Kolmogorov–Smirnov test: p= 0.14, Mann–Whitney U test: p= 0.030; FX naïve: 24 cells, FX post: 22 cells).

**(G)** Same as (F) but for IPSCs. (Kolmogorov–Smirnov test: p= 0.92, Mann–Whitney U test: p= 0.94; WT naïve: 15 cells, WT post: 17 cells).

**(H)** Same as (F) but for IPSCs. (Kolmogorov–Smirnov test: p= 0.17, Mann–Whitney U test: p= 0.034; WT naïve: 24 cells, WT post: 22 cells).

**(I)** same as (E) but from L5 PCs. (Kolmogorov–Smirnov test: p= 0.060, Mann–Whitney U test: p= 0.025; WT naïve: 14 cells, WT post: 15 cells).

**(J)** Same as (I) but from FX pre and post-training mice groups. (Kolmogorov–Smirnov test: p= 0.58, Mann–Whitney U test: p= 0.70; WT naïve: 17 cells; WT post: 22 cells).

**(K)** Same as (I) but for IPSCs. (Kolmogorov–Smirnov test: p= 0.95, Mann–Whitney U test: p= 1.0; WT naïve: 16 cells; WT post: 16 cells).

**(M)** Same as (J) but for IPSCs. (Kolmogorov–Smirnov test: p= 0.70, Mann–Whitney U test: p= 0.62; WT naïve: 17 cells; WT post: 22 cells).

Error bars indicate SEM. $*p < 0.05$; $**p < 0.01$; $***p < 0.001$.

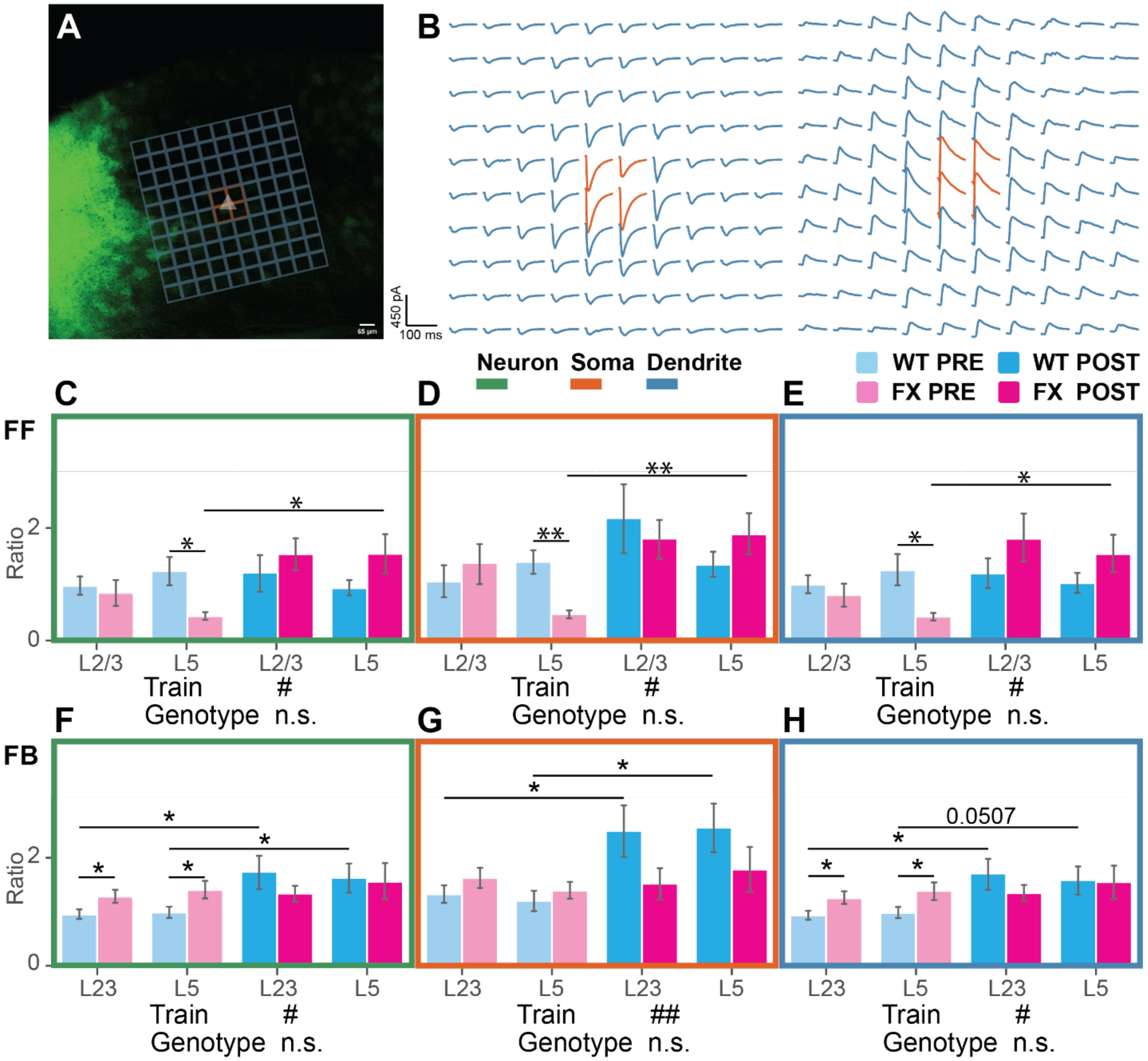


**Figure S9. E/I ratio of FF and FB eCRACM and iCRACM in LM<->V1 reciprocal pathway, related to Figure 4 and 5.**

**(A)** Examples of a confocal image of a coronal slice overlaid by the grids of CRACM patterns. The white triangle indicates where the cell body should be. The area overlaid by orange grids was considered soma areas, while blue grids are considered dendric regions.

**(B)** Examples of $eCRACM_{EPSC}$ traces (left) and $iCRACM_{IPSC}$ traces (right). Orange traces are from soma areas, and blue traces are from dendrite areas.

**(C)** Bar graph of averaged E/I ratio from each grid of CRACM map for whole neuron areas in all groups of mice (WT pre: N=24, WT post: N=36, FX pre: N=25, FX post: N=37 cells). Significance between perceptual experience and genotype was reported from two-way ANOVA (genotype: F= 0.266, p= 0.607; perceptual experience: F= 4.90, p= 0.0288). In either L2/3 or L5, significance was reported by two-way ANOVA followed by Tukey's HSD tests (L2/3: two-way ANOVA: genotype: F = 0.236; p = 0.629. Perceptual experience: F = 2.59; p = 0.113. Tukey's post hoc: WT pre versus FX pre: p = 0.69; WT post versus FX post:

p = 0.46; WT pre versus WT post: p = 0.62; FX pre versus FX post: p = 0.091. L5: two-way ANOVA: genotype: F = 0.032; p = 0.86. Perceptual experience: F = 2.23; p = 0.141. Tukey's post hoc: WT pre versus FX pre: p = 0.011; WT post versus FX post: p = 0.106; WT pre versus WT post: p = 0.29; FX pre versus FX post: p = 0.016).

**(D)** Same as (C) but for the soma area. Significance between train and genotype was reported from two-way ANOVA (genotype: F= 1.24, p= 0.267; perceptual experience: F= 6.10, p= 0.0149). In either L2/3 or L5, significance was reported by two-way ANOVA followed by Tukey's HSD tests (L2/3: two-way ANOVA: genotype: F = 1.18; p = 0.28. Perceptual experience: F = 3.91; p = 0.053. Tukey's post hoc: WT pre versus FX pre: p = 0.536; WT post versus FX post: p = 0.294; WT pre versus WT post: p = 0.28; FX pre versus FX post: p = 0.11. L5: two-way ANOVA: genotype: F = 0.034; p = 0.855. Perceptual experience: F = 5.51; p = 0.0224. Tukey's post hoc: WT pre versus FX pre: p = 1E-3; WT post versus FX post: p = 0.223; WT pre versus WT post: p = 0.886; FX pre versus FX post: p = 5.1E-3).

**(E)** Same as (C) but for the dendritic area. Significance between train and genotype was reported from two-way ANOVA (genotype: F= 0.256, p= 0.613; perceptual experience: F= 4.32, p= 0.040). In either L2/3 or L5, significance was reported by two-way ANOVA followed by Tukey's HSD tests (L2/3: two-way ANOVA: genotype: F = 0.208; p = 0.65. Perceptual experience: F = 2.37; p = 0.129. Tukey's post hoc: WT pre versus FX pre: p = 0.52; WT post versus FX post: p = 0.41; WT pre versus WT post: p = 0.74; FX pre versus FX post: p = 0.083. L5: two-way ANOVA: genotype: F = 0.032; p = 0.86. Perceptual experience: F = 2.23; p = 0.141. Tukey's post hoc: WT pre versus FX pre: p = 0.013; WT post versus FX post: p = 0.092; WT pre versus WT post: p = 0.23; FX pre versus FX post: p = 0.018).

**(F)** Bar graph of averaged E/I ratio from each grid of CRACM map for whole neuron areas in all groups of mice (WT pre: N=32, WT post: N=35, FX pre: N=35, FX post: N=42 cells). Significance between perceptual experience and genotype was reported from two-way ANOVA (genotype: F= 0.106, p= 0.745; perceptual experience: F= 6.227, p= 0.0137). In either L2/3 or L5, significance was reported by two-way ANOVA followed by Tukey's HSD tests (L2/3: two-way ANOVA: genotype: F = 0.0893; p = 0.766. Perceptual experience: F = 4.247; p = 0.0429. Tukey's post hoc: WT pre versus FX pre: p = 0.0468; WT post versus FX post: p = 0.247; WT pre versus WT post: p = 0.0299; FX pre versus FX post: p = 0.777.

L5: two-way ANOVA: genotype: F = 0.441; p = 0.509. Perceptual experience: F = 2.109; p = 0.151. Tukey's post hoc: WT pre versus FX pre: p = 0.0493; WT post versus FX post: p = 0.871; WT pre versus WT post: p = 0.0434; FX pre versus FX post: p = 0.710).

**(G)** Same as (C) but for the soma area. Significance between train and genotype was reported from two-way ANOVA (genotype: F= 1.942, p= 0.166; perceptual experience: F= 7.28, p= 0.00786). In either L2/3 or L5, significance was reported by two-way ANOVA followed by Tukey's HSD tests (L2/3: two-way ANOVA: genotype: F = 1.407; p = 0.239. Perceptual experience: F = 2.52; p = 0.117. Tukey's post hoc: WT pre versus FX pre: p = 0.240; WT post versus FX post: p = 0.0847; WT pre versus WT post: p = 0.0331; FX pre versus FX post: p = 0.771.
L5: two-way ANOVA: genotype: F = 0.596; p = 0.443. Perceptual experience: F = 4.684; p = 0.0343. Tukey's post hoc: WT pre versus FX pre: p = 0.471; WT post versus FX post: p = 0.243; WT pre versus WT post: p = 0.0198; FX pre versus FX post: p = 0.484).

**(H)** Same as (C) but for the dendritic area. Significance between train and genotype was reported from two-way ANOVA (genotype: F= 0.193, p= 0.661; perceptual experience: F=

6.581, p= 0.0113). In either L2/3 or L5, significance was reported by two-way ANOVA followed by Tukey's HSD tests (L2/3: two-way ANOVA: genotype: F = 0.0462; p = 0.830. Perceptual experience: F = 4.736; p = 0.0328. Tukey's post hoc: WT pre versus FX pre: p = 0.0424; WT post versus FX post: p = 0.290; WT pre versus WT post: p = 0.0304; FX pre versus FX post: p = 0.627.

L5: two-way ANOVA: genotype: F = 0.547; p = 0.462. Perceptual experience: F = 2.092; p = 0.153. Tukey's post hoc: WT pre versus FX pre: p = 0.0467; WT post versus FX post: p = 0.934; WT pre versus WT post: p = 0.0507; FX pre versus FX post: p = 0.684).

Data were presented as mean ± SEM. Two-way ANOVA: #p<0.05, ##p<0.01, ###p<0.001, ####p<0. 0001.Tukey HSD' test: *p<0.05, **p<0.01, ***p<0.001, ****p<0.0001

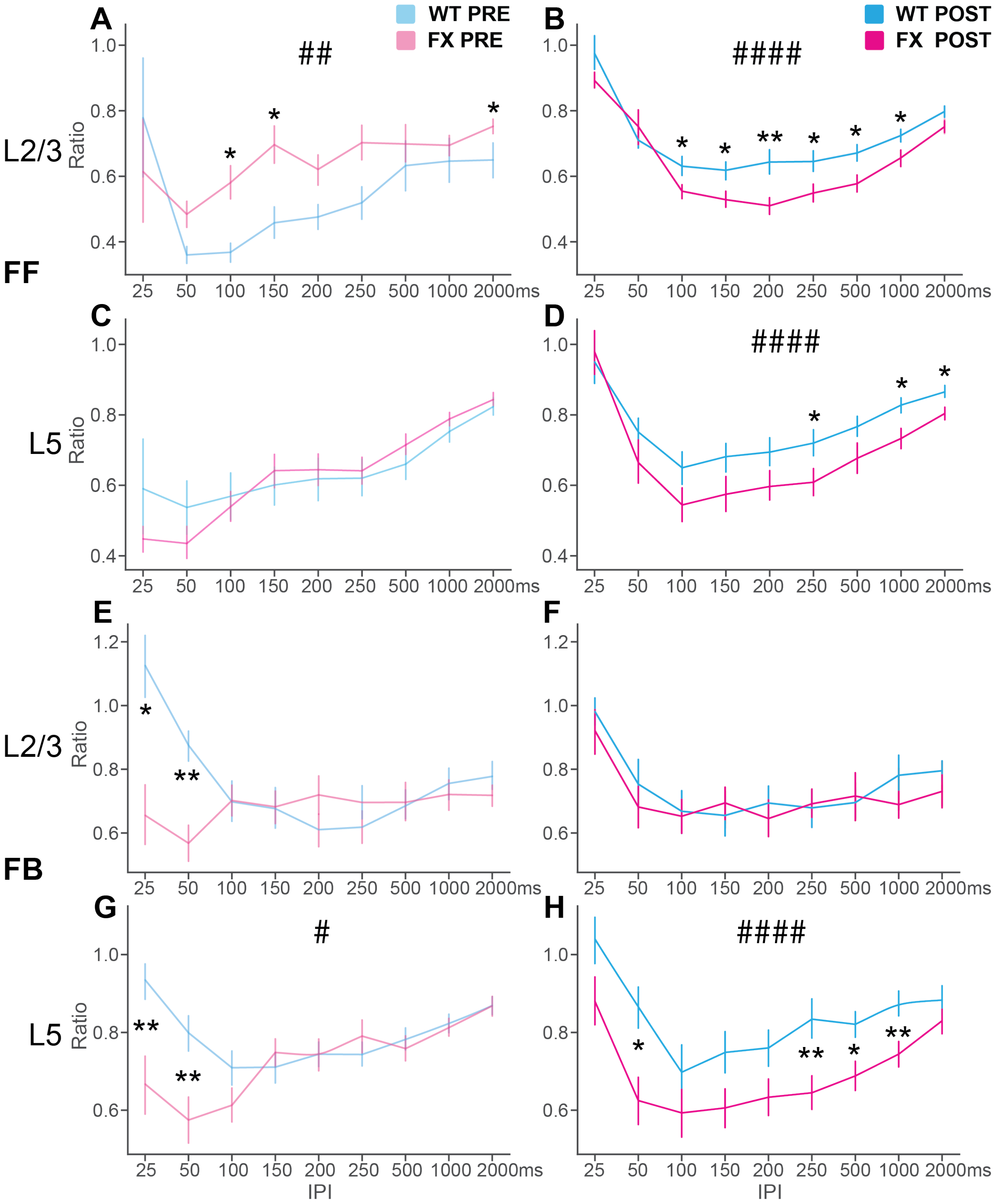


**Figure S10. Paired pulse facilitation of FF V1 to LM and FB LM to V1 L2/3 and L5 PCs projections comparing genotypes, related to Figure 6.**

**(A)** Average PPRs of naïve WT and FX mice of corresponding inter-pulse interval in $FF_{V1\text{->}LM\ L2/3}$ pathway. Significance was reported from two-way ANOVA (perceptual experience: F = 9.70;

p = 2.104E-3) and Tukey's HSD tests (25ms, p= 0.535, WT: N=7, FX: N=11 cells; 50 ms, p= 0.222, WT: N=3, FX: N=15 cells; 100 ms p= 0.0261, WT: N=6, FX: N=17 cells, 150 ms, p= 0.0211, WT: N=7, FX: N=17 cells, 200 ms, p= 0.0561, WT: N=15, FX: N=19 cells, 250 ms, p= 0.0576, WT: N=15, FX: N=18 cells, 500 ms, p= 0.5396, WT: N=8, FX: N=20 cells, 1000 ms p= 0.4684, WT: N=8, FX: N=20 cells; 2000 ms p= 0.0479, WT: N=15, FX: N=21 cells).

**(B)** Same as (A), but from post-training groups. Significance was reported from two-way ANOVA (perceptual experience: F = 27.38; p = 2.90E-07) and Tukey's HSD tests (25ms, p= 0.3283, WT: N=16, FX: N=7 cells; 50 ms, p= 0.4861, WT: N=18, FX: N=16 cells; 100 ms p= 0.0434, WT: N=20, FX: N=19 cells, 150 ms, p= 0.022, WT: N=21, FX: N=17 cells, 200 ms, p= 0.0095, WT: N=23, FX: N=21 cells, 250 ms, p= 0.0288, WT: N=23, FX: N=22 cells, 500 ms, p= 0.0137, WT: N=24, FX: N=22 cells, 1000 ms, p= 0.0386, WT: N=25, FX: N=21 cells; 2000 ms p= 0.0817, WT: N=25, FX: N=21 cells).

**(C)** Average PPRs of naïve WT and FX mice of corresponding inter-pulse intervals in $FF_{V1\text{->}LM\ L5}$ pathway. Significance was reported from two-way ANOVA (perceptual experience: F = 0.0325; p = 0.857) and Tukey's HSD tests (25ms, p= 0.2661, WT: N=9, FX: N=12 cells; 50 ms, p= 0.2442, WT: N=11, FX: N=15 cells; 100 ms, p= 0.721, WT: N=24, FX: N=20 cells, 150 ms, p= 0.6064, WT: N=24, FX: N=21 cells, 200 ms, p= 0.7534, WT: N=11, FX: N=16 cells, 250 ms, p= 0.7593, WT: N=11, FX: N=17 cells, 500 ms, p= 0.3026, WT: N=11, FX: N=18 cells, 1000 ms, p= 0.334, WT: N=11, FX: N=18 cells; 2000 ms, p= 0.5449, WT: N=11, FX: N=18 cells).

**(D)** Same as (C), but from post-training groups. Significance was reported from two-way ANOVA (perceptual experience: F = 18.35; p = 2.340E-05) and Tukey's HSD tests (25ms, p= 0.7664, WT: N=18, FX: N=16 cells; 50 ms, p= 0.2472, WT: N=23, FX: N=14 cells; 100 ms p= 0.1406, WT: N=20, FX: N=19 cells, 150 ms, p= 0.1061, WT: N=24, FX: N=21 cells, 200 ms, p= 0.1066, WT: N=24, FX: N=23 cells, 250 ms, p= 0.0472, WT: N=24, FX: N=23 cells, 500 ms, p= 0.0867, WT: N=24, FX: N=21 cells, 1000 ms, p= 0.0114, WT: N=24, FX: N=22 cells; 2000 ms p= 0.0151, WT: N=24, FX: N=23 cells).

**(E)** Average PPRs of naïve WT and FX mice of corresponding inter-pulse intervals in $FB_{LM\text{->}V1\ L2/3}$ pathway. Significance was reported from two-way ANOVA (perceptual experience: F = 1.34; p = 0.248) and Tukey's HSD tests (25 ms, p= 0.030, WT: N=5, FX: N=21 cells; 50 ms, p= 0.002, WT: N=9, FX: N=19 cells; 100 ms p= 0.9, WT: N=15, FX: N=26 cells, 150 ms, p= 0.9, WT: N=16, FX: N=23 cells, 200 ms, p= 0.250, WT: N=16, FX: N=27 cells, 250 ms, p= 0.350, WT: N=15, FX: N=25 cells, 500 ms, p= 0.9, WT: N=16, FX: N=26 cells, 1000 ms p= 0.664, WT: N=17, FX: N=27 cells; 2000 ms p= 0.272, WT: N=17, FX: N=27 cells).

**(F)** Same as (E), but from post-training groups. Significance was reported from two-way ANOVA (perceptual experience: F = 1.066; p = 0.303) and Tukey's HSD tests (25ms, p= 0.597, WT: N=7, FX: N=14 cells; 50 ms, p= 0.528, WT: N=10, FX: N=20 cells; 100 ms p= 0.868, WT: N=12, FX: N=20 cells, 150 ms, p= 0.658, WT: N=13, FX: N=21 cells, 200 ms, p= 0.592, WT: N14, FX: N=25 cells, 250 ms, p= 0.869, WT: N=13, FX: N=25 cells, 500 ms, p= 0.854, WT: N=16, FX: N=29 cells, 1000 ms, p= 0.209, WT: N=17, FX: N=29 cells; 2000 ms p= 0.387, WT: N=31, FX: N=17 cells).

**(G)** Average PPRs of naïve WT and FX mice of corresponding inter-pulse intervals $FB_{LM\text{->}V1\ L5}$ pathway. Significance was reported from two-way ANOVA (perceptual experience: F = 6.55; p = 0.011) and Tukey's HSD tests (25ms, p= 8.4E-3, WT: N=19, FX: N=24 cells; 50 ms, p= 9.1E-3, WT: N=21, FX: N=28 cells; 100 ms p= 0.132, WT: N=27, FX: N=31 cells, 150 ms, p= 0.486, WT: N=30, FX: N=30 cells, 200 ms, p= 0.9, WT: N=30, FX: N=31 cells, 250 ms, p=

0.370, WT: N=32, FX: N=31 cells, 500 ms, p= 0.604, WT: N=30, FX: N=31 cells, 1000 ms, p= 0.780, WT: N=30, FX: N=31 cells; 2000 ms p= 0.9, WT: N=29, FX: N=30 cells).

**(H)** Same as (G), but from post-training groups. Significance was reported from two-way ANOVA (perceptual experience: F = 34.69; p = 8.925E-9) and Tukey's HSD tests (25ms, p= 0.1004, WT: N=11, FX: N=19 cells; 50 ms, p= 0.0115, WT: N=15, FX: N=23 cells; 100 ms p= 0.273, WT: N=18, FX: N=23 cells, 150 ms, p= 0.0802, WT: N=18, FX: N=25 cells, 200 ms, p= 0.079, WT: N=19, FX: N=24 cells, 250 ms, p= 0.007, WT: N=20, FX: N=25 cells, 500 ms, p= 0.0144, WT: N=20, FX: N=25 cells, 1000 ms, p= 8.3E-3, WT: N=20, FX: N=25 cells; 2000 ms p= 0.290, WT: N=20, FX: N=25 cells).

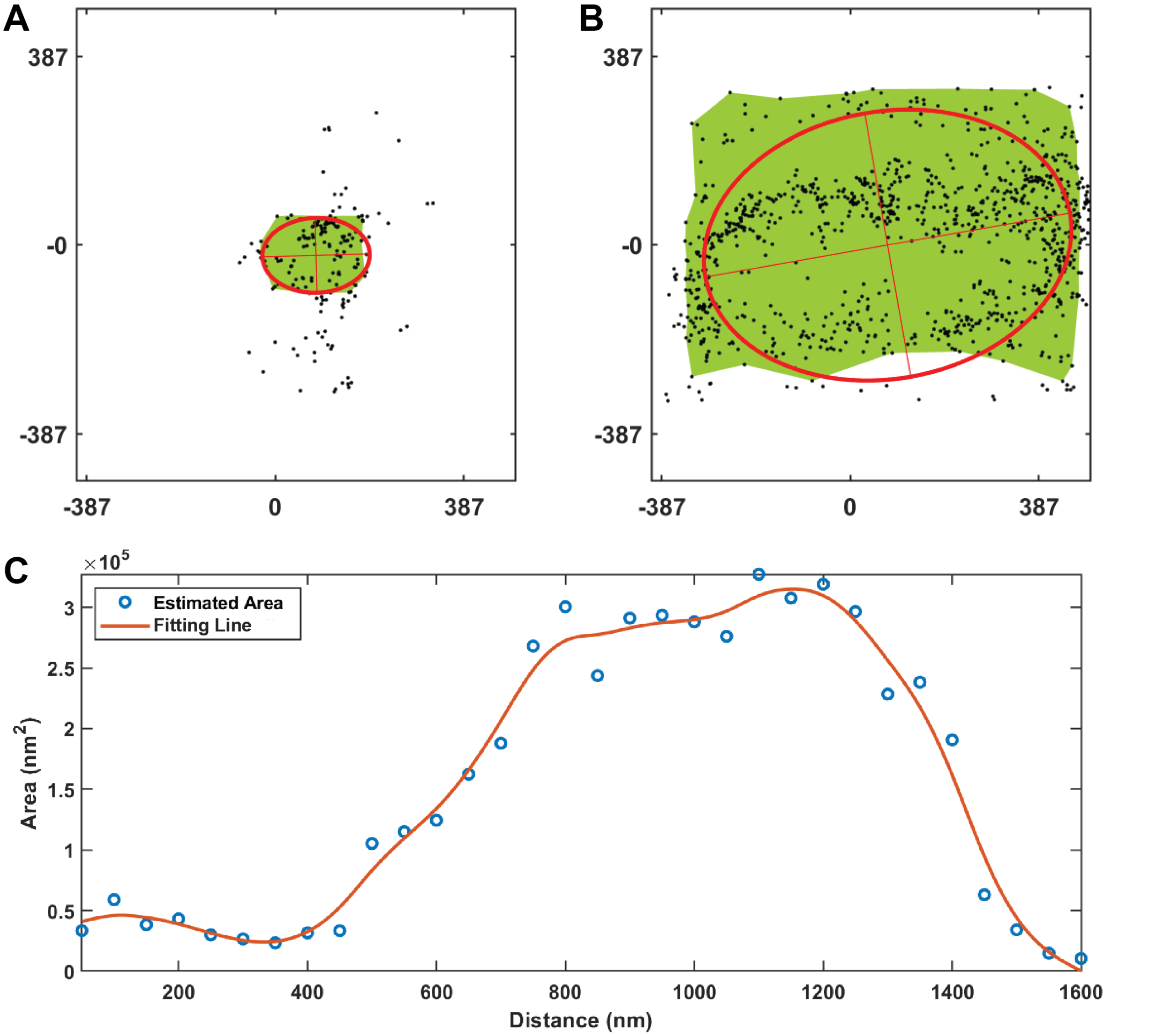


**Figure S11. Morphology analysis of dendritic spines from apical dendrites of L5 PCs, related to Figure 7.**

**(A)** Acquiring the size of cross-sections of the neck by fitting the scattered point contour with Alpha-shape in Figure 8C top.

**(B)** same as (A) but for the head of spine Figure 9C bottom.
**(C)** Plot of the area of the cross-section along the axial of the spine.
